# SARS-CoV-2 spike opening dynamics and energetics reveal the individual roles of glycans and their collective impact

**DOI:** 10.1101/2021.08.12.456168

**Authors:** Yui Tik Pang, Atanu Acharya, Diane L. Lynch, Anna Pavlova, James C. Gumbart

**Affiliations:** School of Physics, Georgia Institute of Technology, Atlanta, GA 30332

**Author notes:** These authors contributed equally to this work (Yui Tik Pang, Atanu Acharya). BioInspired Syracuse and Department of Chemistry, Syracuse University, Syracuse, NY 13244.

## Abstract

The trimeric spike (S) glycoprotein, which protrudes from the SARS-CoV-2 viral envelope, binds to human ACE2, initiated by at least one protomer’s receptor binding domain (RBD) switching from a “down” (closed) to an “up” (open) state. Here, we used large-scale molecular dynamics simulations and two-dimensional replica exchange umbrella sampling calculations with more than a thousand windows and an aggregate total of 160 µs of simulation to investigate this transition with and without glycans. We find that the glycosylated spike has a higher barrier to opening and also energetically favors the down state over the up state. Analysis of the S-protein opening pathway reveals that glycans at N165 and N122 interfere with hydrogen bonds between the RBD and the N-terminal domain in the up state, while glycans at N165 and N343 can stabilize both the down and up states. Finally we estimate how epitope exposure for several known antibodies changes along the opening path. We find that the BD-368-2 antibody’s epitope is continuously exposed, explaining its high efficacy.

## Introduction

The COVID-19 pandemic caused by the SARS-CoV-2 coronavirus quickly spread worldwide with unprecedented detrimental impact on global health and economies^1^. The rapid development of several vaccines^2^, monoclonal antibody treatments^3^, and therapeutics^4^ have mitigated the current viral outbreak. However, the ongoing threat of variants, including the now dominant Omicron sub-lineages^5^, and the possibility of future coronavirus outbreaks^6^ necessitate a thorough understanding of the viral life cycle, including recognition, binding, infection, and immune response.

SARS-CoV-2 infection is initiated by the recognition of, and binding to, the host-cell angiotensin-converting enzyme 2 (ACE2) receptor^7,8^. This process is mediated by the SARS-CoV-2 spike (S) protein, a homotrimeric class I fusion glycoprotein that protrudes from the surface of the SARS-CoV-2 virion. Release of the S-protein sequence in early 2020, combined with earlier structural work on related betacoronaviruses, led to the rapid determination of structures of solublized, pre-fusion stabilized S-protein ectodomain constructs^9–11^ (Fig. 1) as well as full length spike^12^ and intact virions^13^. Each protomer consists of the S1 and S2 subunits separated by a multibasic furin cleavage site. S1 contains the receptor binding domain (RBD) and mediates host cell recognition while S2 consists of the membrane fusion machinery necessary for viral entry^7^. The S-protein is a major antigenic target with multiple epitopes that are targeted by the human immune system, including the RBD and the N-terminal domain (NTD)^14–17^. Recombinant RBD is also suggested to be a potential entry inhibitor against SARS-CoV-2^18^. Moreover, glycosylation of the S-protein aids in masking and shielding the virus from host immune system response^19–21^. The S-protein is characterized by down and up conformational states, which transiently interconvert via a hinge-like motion exposing the receptor binding motif (RBM), which is composed of RBD residues S438 to Q506^22^. The RBM is buried in the inter-protomer interface of the down S-protein; therefore, binding to ACE2 relies on the stochastic interconversion between the down and up states.

**Figure 1.**
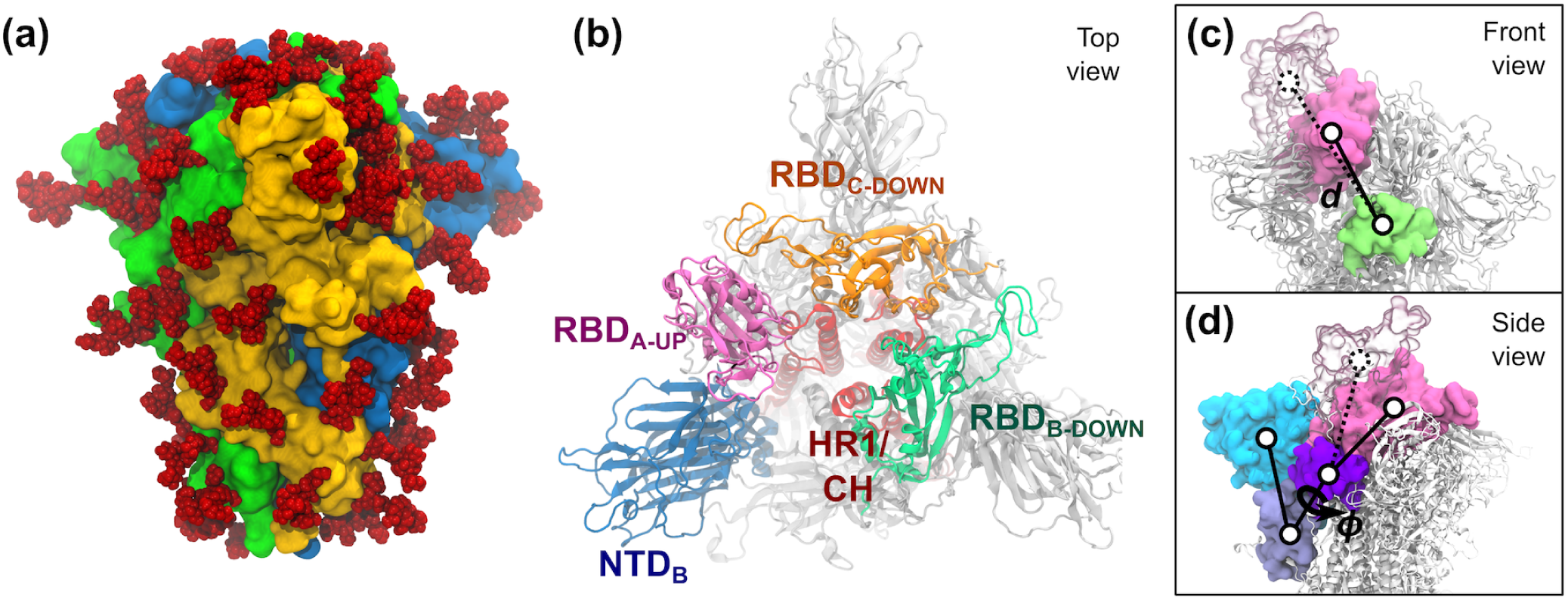
S-protein of SARS-CoV-2. (a) The trimeric S-protein in the all-down state, colored by protomer. Glycans are shown as red spheres. (b) Top view of the S-protein in the one-up state. Important domains of the spike are highlighted, including the N-terminal domain (NTD, 14–306), the receptor binding domains (RBD, 336–518), the heptad repeat 1 (HR1, 908–986), and the central helix (CH, 987–1035). (c,d) The two collective variables defined to describe the opening of RBD-A include: (c) the center-of-mass distance *d* between RBD-A (336–518, pink) and SD1-B (531–592, lime), and (d) the dihedral angle ϕ formed by the center of mass of the domains RBD-A (336–518, pink), SD1-A (531–592, purple), SD2-A (593–677, ice blue), and NTD-A (27–307, cyan). RBD-A in both the down (solid pink) and up (transparent pink) states are shown.

Cryo-electron microscopy (cryo-EM) studies have revealed detailed structural information for both the up and down conformational states^23^. However, relatively few studies have explored the dynamics of these up/down states and interconversion between them. For example, single-molecule FRET has been used to demonstrate the stochastic nature of the S-protein transitions^24^, with reported timescales on the order of milliseconds to seconds. Of note, later single-molecule FRET studies find altered down-up transition kinetics for mutant spike proteins^25^. Molecular dynamics (MD) simulations complement these experimental studies by providing the atomic-level descriptions of intermediate states between down and up that are necessary to characterize S-protein opening dynamics. MD simulations have revealed detailed information about the structural stability and the role of glycosylation for both the down and up states, as well as for inter-residue interactions and details of binding to ACE2^20,21,26–29^. Opening pathways determined using steered MD and targeted MD have been reported^29–31^. Additionally, extensive simulations using enhanced sampling techniques such as weighted ensemble^32^ and fluctuation amplification of specific traits (FAST) adaptive sampling combined with Folding@home^33^ have provided details of multiple pathways for the S-protein opening. Moreover, features of the energy landscape of these conformational transitions that are necessary for viral binding and entry are beginning to emerge^27,30,31,34^, including the combination of cryo-EM data with MD simulations to reveal multiple sub-populations for both the down and up states^35,36^.

A recent study by Amaro and coworkers has highlighted the functional role of glycans at N165 and N234 beyond shielding^26^ based on separate equilibrium simulations of the S-protein down and up states. When the RBD transitions to the up state, the glycan at N234 rotates into the resulting void, stabilizing the up conformation. Moreover, MD simulations and mutagenesis have revealed contributions of the glycan at N343 to the dynamics of RBD opening and ACE2 binding^32^. These results suggested a role for the glycan at N343 in the opening conformational transition via “lifting” the RBD through sequential interactions with multiple RBD residues, referred to as “glycan gating”.

Here, we describe newly determined two-dimensional (2D) free-energy landscapes of the SARS-CoV-2 Sprotein opening and closing transitions using replica exchange umbrella sampling (REUS) simulations run on the pre-exascale supercomputer Summit at Oak Ridge National Lab (ORNL) for glycosylated as well as unglycosylated S-protein. We highlight the impact of glycans on each state and on the kinetics of spike opening. Furthermore, we analyzed the exposure of prominent epitopes on the S-protein surface and provide a dynamic picture of antibody binding along the spike-opening path. Finally we report the results of equilibrium MD simulations of the glycosylated and un-glycosylated systems for the down as well as up conformational states in order to further characterize the stabilizing role of the glycans.

## Results

### Glycans modulate the energetics and pathway of spike opening

Using REUS simulations, we studied the free-energy change from the down state of spike to the up state when fully glycosylated as well as when un-glycosylated. We modeled the wild-type (WT) up state based on the diproline mutant structure from Walls et al.^10^ (PDB: 6VYB). The down-state was modeled using a more recent structure from Cai et al.^12^ (PDB: 6XR8) without the diproline mutations. The glycan with highest population in the mass spectroscopy data from Crispin and coworkers^19^ at each site was added using the GLYCAM Web server developed by the Woods group (http://glycam.org)^37,38^. Additional details of the system are provided in Methods.

We first ran metadynamics simulations of the cryo-EM structures in the down and up states, allowing them to explore the conformational space around them and connect the two structures. The conformational space is described by two collective variables, which are (1) the center-of-mass distance (*d*) between the opening RBD-A and a stationary part of the spike acting as a pivot point, namely the neighboring subdomain 2 on chain B (SD2-B), and (2) the dihedral angle (*ϕ*) formed by the opening RBD-A and other stationary domains on the same protomer, namely SD2-A, SD1-A and NTD-A (Fig. 1c,d). The stationary domains were verified to have minimal dynamics compared to the range of motion of RBD-A (Table S1). Snapshots were then extracted from the metadynamics simulations to seed the REUS simulations, which were run along the same two collective variables. For the glycosylated system, we further performed simulated annealing on the glycans to randomize their conformations in each window. A series of REUS simulations were run to further explore different regions of the *d*-*ϕ* space, using up to 1049 and 1211 windows, and a total simulation time of 65 μs and 91 μs for the glycosylated and un-glycosylated systems, respectively.

For the glycosylated system, the conformations from the two cryo-EM structures emerged as energy minima on the 2D potential of mean force (PMF) as expected (Fig. 2a). Between the two stable conformations, the up state possesses a higher energy than the down state by 5.2 ± 0.1 kcal/mol (Figs. 2c, S1a). This finding is consistent with multiple computational studies using various enhanced sampling methods, which also concluded that the down state is the more probable conformation^33,34,39,40^. The differences between the two end-state energy wells does not end with their depth but also their breadth. If we measure the width of the energy wells along *d* at 2.5 kcal/mol above their energy minima, the down-state energy well spans from *d* = 43.1 Å to 48.9 Å, while the up-state energy well covers *d* = 60.1 Å to 75.6 Å, making the latter 2.7 times the width of the former. Recent works from Zimmerman et al.^33^ and Sztain et al.^32^ also find the RBD to be highly flexible in the up state, allowing a wide range of up-state structures that open wider than the cryo-EM structure. Our PMF now allows one to estimate the likelihood of observing such widely open structures. We also computed the minimum energy path (MEP)^41,42^ connecting the down and up states (Fig. 2a). The path is mostly diagonal on the 2D PMF, with the exceptions of a sharp increase of *ϕ* while exiting the down-state energy well and a slight decrease of *ϕ* when entering the up-state energy well. The energy barrier separating the two stable conformations has a height of 11.0 ± 0.1 kcal/mol and is located at *d* = 55.6 Å.

**Figure 2.**
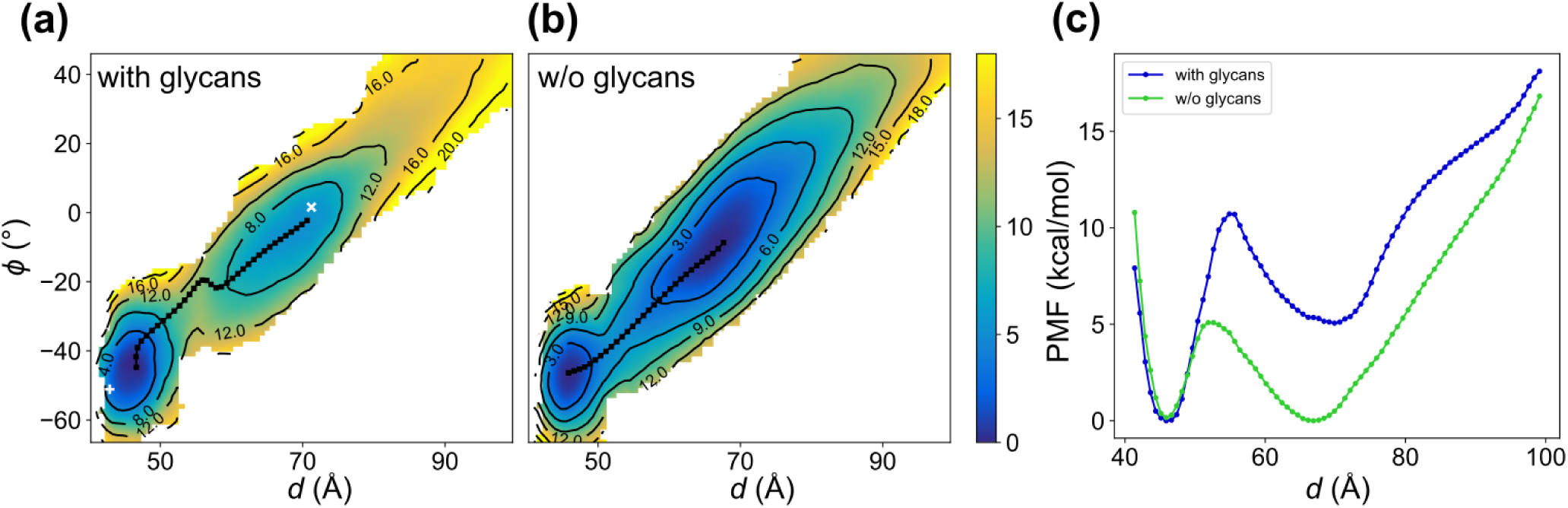
PMFs describing the opening of RBD-A. (a,b) The 2D PMFs of the (a) glycosylated and (b) un-glycosylated systems along two collective variables, *d* and *ϕ*, defined to describe the opening of RBD-A (Fig. 1c,d). The location of the down(6XR8) and up-state (6VYB) cryo-EM structures are indicated in (a) with a “+” and “x” sign, respectively. The black dotted line shows the MEP for each system. (c) The free energies are projected onto *d* and plotted as 1D PMFs.

The PMF for the un-glycosylated system is qualitatively similar but significantly different quantitatively from the glycosylated one (Fig. 2b). Without the glycans, the energy difference between the down and up states becomes surprisingly small and falls below the statistical error (Figs. 2c, S1b). The width ratio along *d* between the down- and up-state energy wells remains at 2.7, the same as the glycosylated system, with the down-state well spanning from *d* = 43.6 Å to 49.1 Å and the up-state well from *d* = 59.0 Å to 74.1 Å. The much wider energy well in the up state results in a dominating equilibrium population of 88% over the down-state population of 12%, despite the small energy difference between them. Both the position of the up-state energy minimum and the energy barrier shifting towards the down state, moving from *d* = 70.6 Å to 67.6 Å and from *d* = 55.6 Å to 52.6 Å, respectively. We also note that the height of energy barrier between the two stable conformations is reduced to 5.1± 0.1 kcal/mol. Combining the above information, our results indicate that the removal of glycans not only flips the downto up-state population ratio but also impacts the extent of the RBD opening at the up-state minimum, both possibly disruptive to the functionality of the S-protein. Experiments have also shown that removing glycans around the RBD (N234, N165, and N343)^26,32^ or decreasing the glycan complexity^43^ leads to a decrease in ACE2 binding.

### Kinetics of opening and binding are altered by the removal of glycans

To quantify the kinetics of S-protein opening, we computed the mean first passage time (MFPT) along the MEP using the Smoluchowski diffusion equation for the glycosylated system^44,45^. The Smoluchowski diffusion equation provides a simple description of a Brownian “particle”, relying only on the diffusion coefficient and free energy along the MEP. Additional constrained simulations were run to determine the diffusion coefficient along the MEP using the velocity autocorrelation function (VACF)^46,47^. The MFPT from the down state to the up state is 1478.5 ms, while the reverse is 0.9 ms. Comparing with experimental observation by single-molecule FRET^24^, it appears that the down-to-up MFPT is overestimated by ∼5× and that for the reverse direction is underestimated by ∼100 ×, possibly due to the fact that we did not account for the long timescale for glycans to equilibrate when computing the diffusion coefficient. We also approximated the MFPT for the un-glycosylated system. The reduced transition barrier in the PMF (Fig. 2c) led to a concomitant reduction in the MFPT, which is 143 μs for the down-to-up transition and 497 μs for the up-to-down transition.

Next, we considered the chemical dynamics according to the following paired reactions:

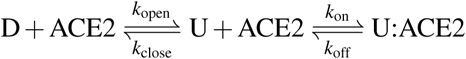

in which D represents the down state of the S-protein, U the up state, and U:ACE2 the bound state (Fig. 3). The open/close rates (*k*_open_/*k*_close_) are determined by the MFPTs and the binding/unbinding rates (*k*_on_/*k*_off_) for RBD alone to ACE2 are taken from Lan et al.^22^. After solving the Master equation for these reactions ([ACE2]_*i*_ = 15 nM; see Supplementary Information), we find that for the un-glycosylated S-protein, the populations are 70% bound, 23% up-but-unbound, and only 7% down (Fig. 3b). Deep mutational scanning of the RBD found any mutation to N343, which eliminates the glycan bound at this position, is detrimental, with an inferred ΔΔ*G* of +0.35 to 2.03 kcal/mol^48^. Using our approximate open/close rates for the un-glycosylated S-protein but modifying the binding/unbinding rates according to the mutation data, we find that the bound-state population decreases to between 8% (ΔΔ*G* = 2.03 kcal/mol) and 57% (ΔΔ*G* = 0.35 kcal/mol). Thus, even though removing all glycans increases the favorability of the up state, an associated reduction in binding affinity could eliminate the otherwise expected gain in the ACE2-bound population.

**Figure 3.**
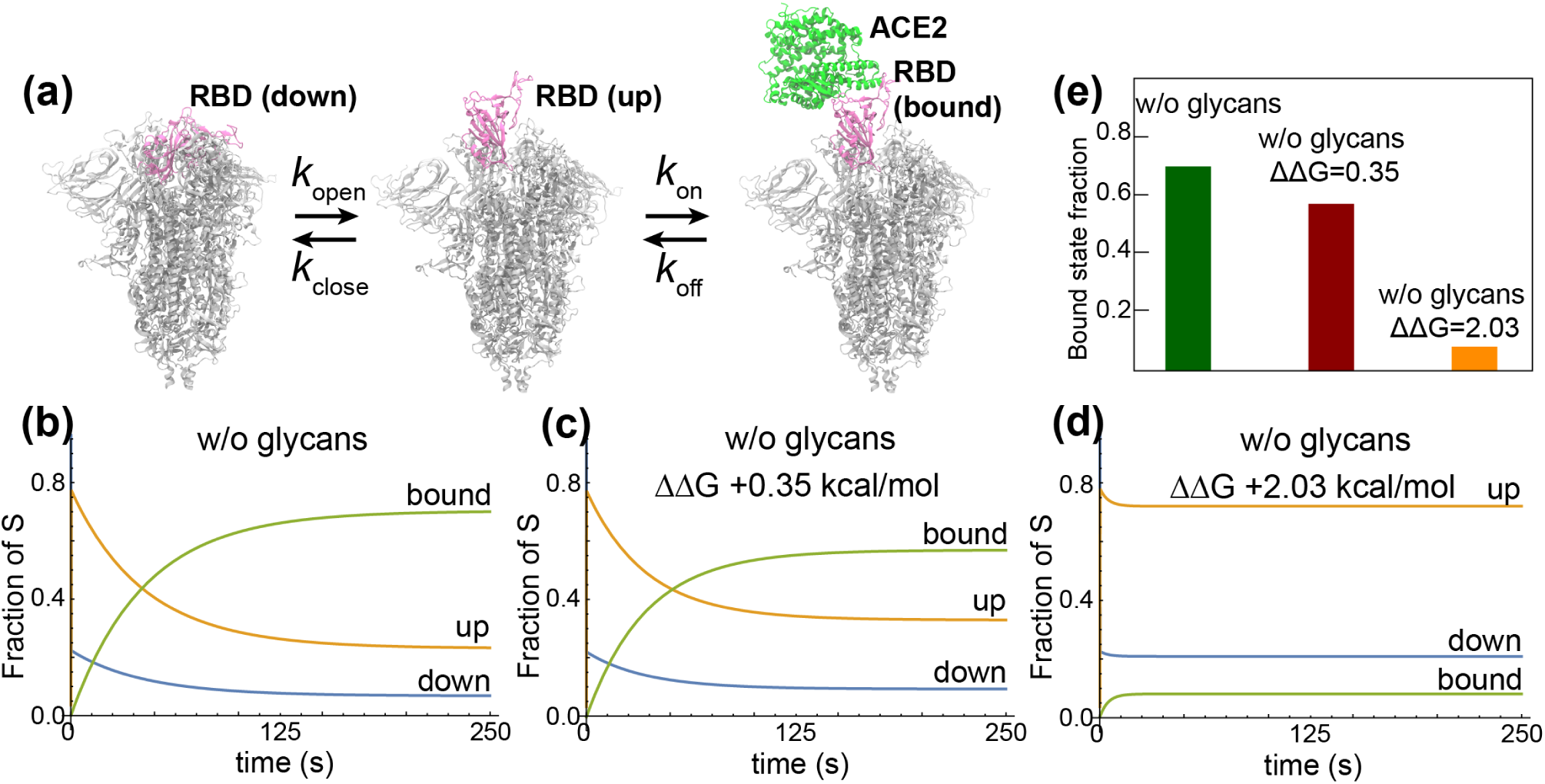
Kinetics of S-protein opening and closing. (a) Transitions between RBD-down, up, and bound states are shown with their associated rates. (b-d) Fraction of S proteins in each state (up, down, and ACE2-bound) under different conditions, namely (b) with no glycans, (c) with no glycans and an assumed increase in the free energy of binding of RBD to ACE2 of 0.35 kcal/mol, and (d) with no glycans and an increase in binding free energy of 2.03 kcal/mol. (e) Bound-state fraction at equilibrium for the three conditions in (b-d).

### Glycans interfere with hydrogen bonds that stabilize the up-state RBD

To better characterize the effects of glycans on the energetics of spike opening, we examined how the RBD interacts with the rest of the protein when glycans are present or absent in the REUS simulations. We calculated the number of hydrogen bonds between the opening RBD-A and the rest of the protein and plotted it against the path parameter. (Figs. 4a-b, S2) We found that this number decreases generally as RBD-A opens up, with a distinct increase in interactions after crossing the energy barrier, more so with glycans than without. The total number of hydrogen bonds formed with RBD-A is higher for the glycosylated system, especially in the down state. However, if we only account for protein-protein hydrogen bonds, the un-glycosylated RBD-A was able to form more connections with the other protein domains. The overall declining trend is explained by the fact that RBD-A moves away from the rest of the protein as it opens and consequently breaks contact with the rest of the trimeric S-protein, including neighboring RBDs and S2 domains. However, this movement is insufficient on its own to explain the increase in interactions after crossing the energy barrier as well as the other differences between glycosylated and un-glycosylated simulations.

**Figure 4.**
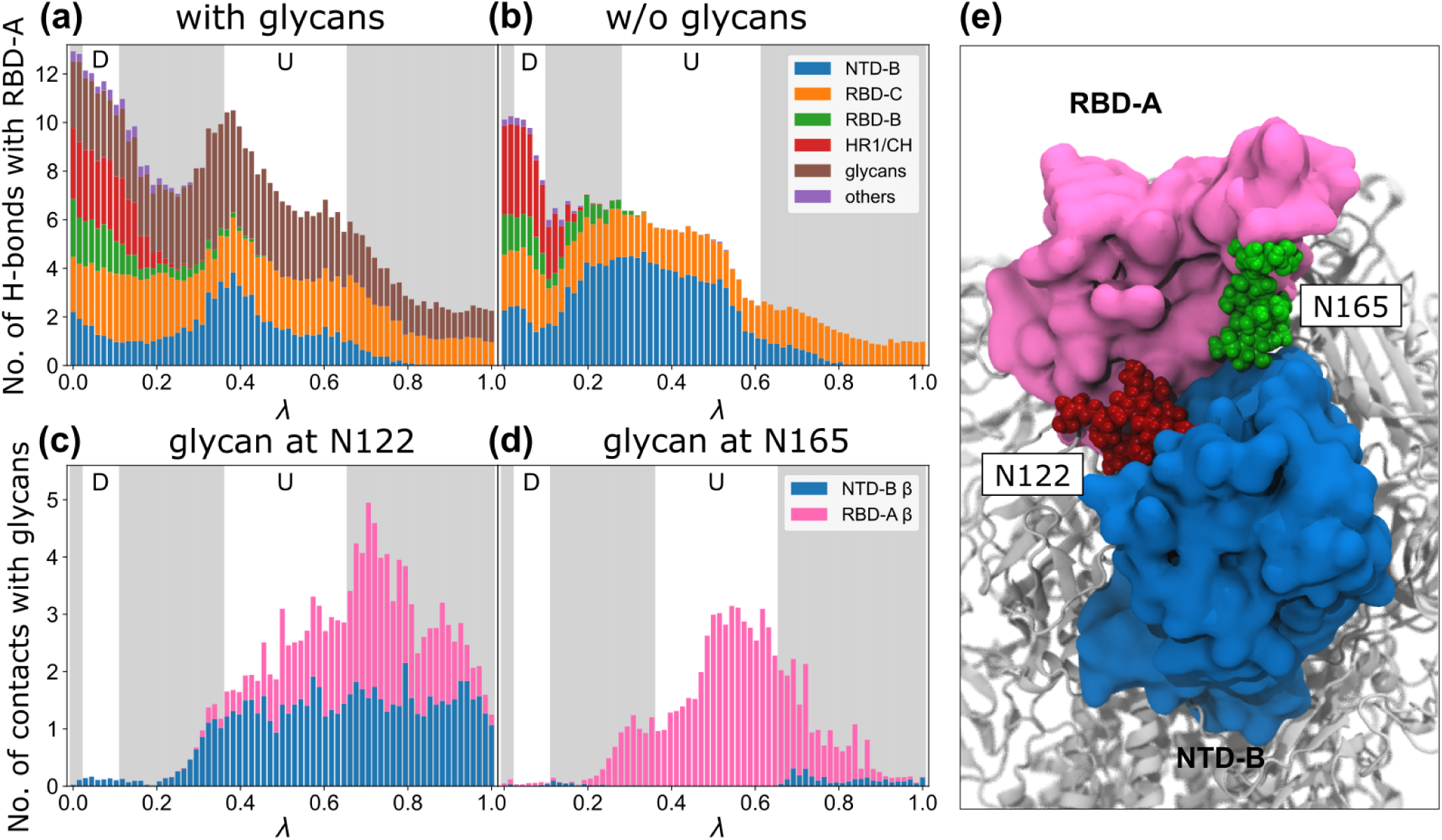
Hydrogen bond analysis reveals the different interaction patterns with and without glycans. (a,b) Along the MEPs as indicated by the path parameter λ (see Fig. S2), the number of hydrogen bonds formed between the opening RBD-A and the rest of the spike are counted and classified by domain for the (a) glycosylated and the (b) un-glycosylated systems. The locations of the down- and up-state energy wells are shown with a white background while other regions are shaded in grey. (c,d) The average number of contacts for the glycosylated system formed between the glycans at (c) N122 and (d) N165 and the neighboring β-strand from NTD-B (165–172) and RBD-A (353–360), which would otherwise form hydrogen bonds with each other if not separated by the glycans. (e) Snapshot from the REUS simulation showing the glycans at N165 and N122 disrupting hydrogen bond formation between RBD-A and NTD-B, destabilizing the up state compared to the un-glycosylated system.

Digging deeper into the hydrogen-bond composition changes along the MEP reveals more about the role of the glycans in the down-to-up state transition. While RBD-A forms a majority of its hydrogen bonds with the other two RBDs and the HR1/CH helix of the S2 unit when it is in the down state, it changes drastically to interacting only with the neighboring NTD-B and RBD-C in the up state (Figs. 4a-b, S3). During this transition, there is a dearth of stabilizing hydrogen bonds, contributing to the energy barrier as seen in the PMFs. With the inclusion of the glycans, they form additional interactions to stabilize the protein, but they also compete with the aforementioned protein-protein interactions. In a compact down-state structure, the existing protein-protein hydrogen bonds are protected from exposure to glycans, leaving the glycans to form new interactions on the protein surface, resulting in an overall increase of the number of hydrogen bonds formed with RBD-A when compared to the un-glycosylated system. However, such protection does not occur in the up state. With RBD-A up and exposed, its interactions with NTD-B are limited. Specifically, most of the hydrogen bonds between RBD-A and NTD-B are displaced by interactions with the glycans at N165 and N122 once RBD-A is lifted high enough for them to intercalate in between (Fig. 4c-e). As a result, the glycans favor the down state indirectly, contributing to the higher free energy of the up state for the glycosylated system.

Extending our hydrogen bond analysis beyond the MEP and to the whole 2D collective variable space furthers our understanding of the differences between the free energy landscapes from the REUS simulations with and without glycans. We plotted the average number of hydrogen bonds between the opening RBD-A and its neighboring NTD-B as a function of the two collective variables, *d* and *ϕ*, showing the increase in interactions between RBD-A and NTD-B is not limited to states along the MEP but also includes a large region on the +*d*-side of the energy barrier (Fig. S4a). Interestingly, the number of hydrogen bonds between RBD-A and NTD-B peaks at *ϕ* = 40° and slowly drops as *ϕ* increases, echoing the previous result that when RBD-A gets further away from the NTD-B, glycans intervene and block interactions between the two domains. On the −*d*-side of the energy barrier, the opposite happens for the hydrogen bonds between RBD-A and RBD-C, where RBD-A and RBD-C are closest to each other when *ϕ* is large and hence has the highest number of hydrogen bonds (Fig. S4b). This observation explains the kink of the MEP when crossing the energy barrier for the glycosylated system. In order to maximize the number of hydrogen bonds stabilizing RBD-A, the spike exits the down-state energy well with a large *ϕ* to maintain maximum contact between RBD-A and RBD-C, before abruptly switching to a smaller *ϕ* when entering the up-state energy well to maximize contact with NTD-B.

### Distinct glycan contacts with RBD-A stabilize both the up and down states

In order to assess the long-timescale dynamics of the glycans, we performed two 2-μs equilibrium simulations of both the glycosylated and un-glycosylated systems. When projected onto the two collective variables used in REUS, all three protomers in the down state for both replicas remained in the down conformation, both with and without glycans (Fig. S5). Although no transition between the up and down states occurred during the equilibrium simulations, the up-states exhibit a much higher flexibility than the down-state ones for both the glycosylated and un-glycosylated systems, reflecting the size difference between the two energy wells as found from the REUS simulations. Interestingly, while the un-glycosylated RBD-A samples around the up-state energy minimum without an obvious directional bias, the glycosylated RBD-A shows a tendency to move towards the −*d* direction from the up-state minimum along the MEP. This result, again, aligns with the REUS simulations showing that the up-state energy well for the glycosylated system has a smaller gradient towards the energy barrier than away from it, whereas the one for the un-glycosylated system is more symmetric (Fig. 2c).

We also analyzed the interactions of glycans around RBD-A, namely those at N165, N234, and N343 of the neighboring chain B, using the equilibrium simulations as well as the REUS trajectories along the MEP. Recently, Amaro and coworkers conducted all-atom MD simulations and mutation experiments to study the individual roles of the above glycans on spike opening^26,32^. Consistent with their results, we observed the swing of the glycan at N234 from pointing outward to inward during spike opening as well as the glycan gating effect by the glycan at N343 (Supplementary Video).

Our simulations also reveal roles for the glycans at N165 and N343 in stabilizing both the up and down states. Specifically, when RBD-A is in the down state, glycans at N165 and N343 wrap around the RBM (Fig. 5a), keeping it in the down state. A similar interaction was observed in another study when a glycan at N370 was artificially introduced; it wrapped around the down-state RBD like a “shoelace”^21^. Remarkably, the glycan at N165 also stabilizes the up-state RBD by supporting part of the exposed RBM (Fig. 5b). The inter-glycan contact between glycans at N343 and N165 is significantly higher in the up state compared to the down state (Fig. S6), suggesting that the glycan at N343 indirectly stabilizes the open state by interacting with the glycan at N165 (Fig. 5b). The two glycans also prevent the flexible RBM from attaching to the neighboring down-state RBD and becoming inaccessible to ACE2, as observed in one of our equilibrium simulations of the un-glycosylated system (Fig. S7).

**Figure 5.**
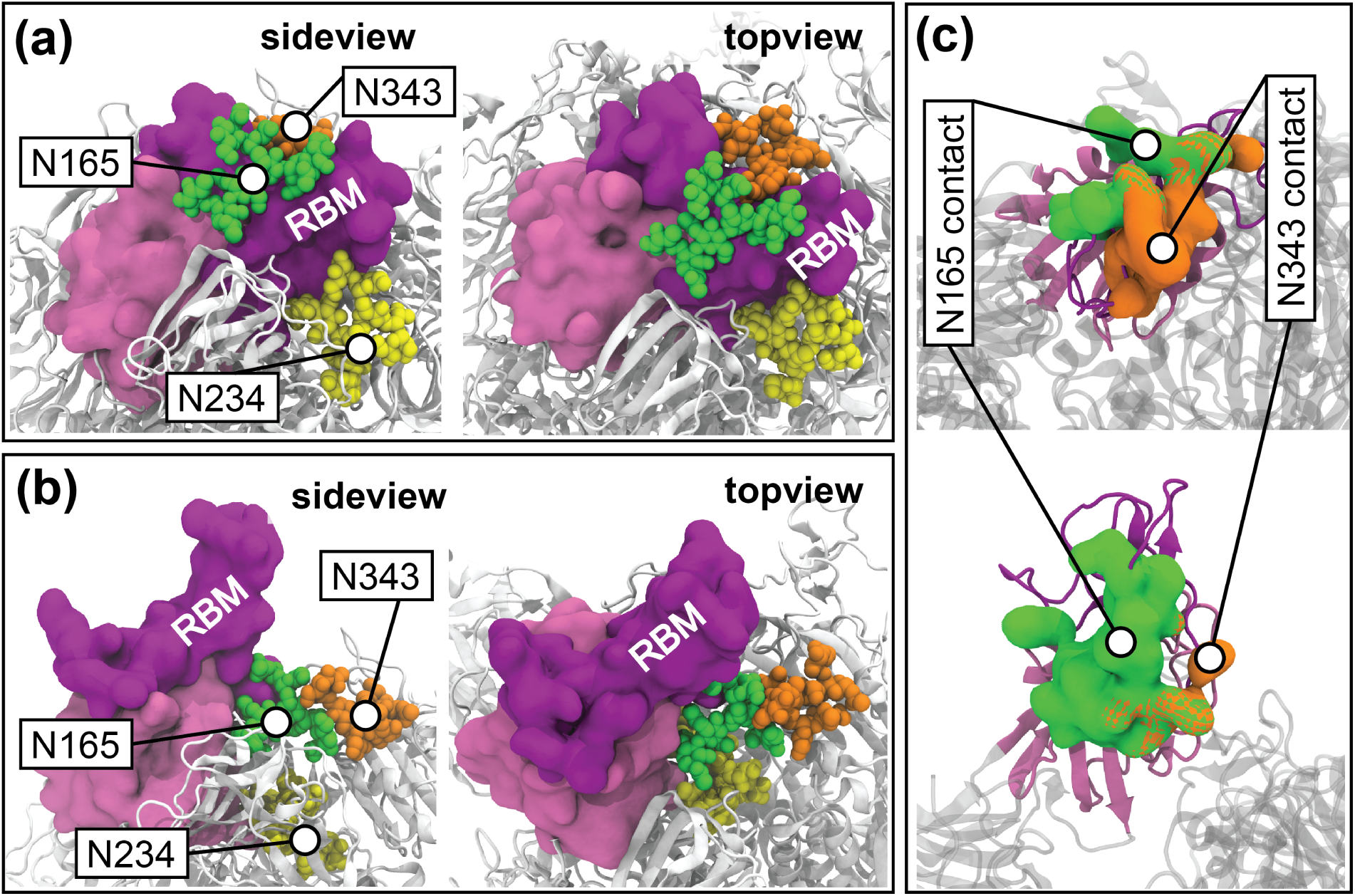
Glycan-protein interactions. Representative locations of S-protein glycans at N165, N234, and N343 in S-protein (a) down and (b) up states extracted from the REUS trajectories. (c) Surface representation of RBD residues in (top) down and (bottom) up states that make contact with glycans at N165 (green) and N343 (orange). The cross-hatch pattern indicates contacts with both glycans. The up and down states of the S-protein were selected from the MEP.

The above observations are confirmed by the contact analysis on the REUS trajectories along the MEP. The glycans at N165 and N343 both interact with the RBM in the down state as illustrated by the average contact values between RBD-A and glycans at N165 and N343 (Figs. S8, S9). As the spike opens, the glycan at N165 switches to interact with RBD-A residues below the RBM, while the glycan at N343 barely contacts RBD-A after the system crosses the activation barrier of opening (Fig. 5c). The contact analysis for the down state also captures the RBM residues (F456, R457, Y489, and F490) involved in glycan gating as previously reported by Sztain et al.^32^.

### Engineered diproline mutations do not affect the equilibrium properties of the ancestral WuhanHu-1 S-protein

S-protein cryo-EM structures often include a double proline mutation, K986P/V987P, located in the turn between HR1 and the CH. Earlier studies on SARS-CoV and MERS-CoV, along with other class I fusion proteins, established that the presence of this proline pair in the HR1-CH turn stabilizes the pre-fusion spike structure and increases protein expression, both important aspects of vaccine design^49^. The S-protein structure from Cai et al.^12^ retains the WT K986/V987 sequence, rather than the double proline mutations commonly made in the production of stabilized pre-fusion spike proteins; they report a shifting inwards (and, thus, tighter packing) of the S1 subunits when compared to the earlier solubilized ectodomain structures^12^. In fact, the loss of a putative salt bridge between K986 and an aspartic acid (D427/D428) on an adjacent protomer has been implicated in producing the less tightly packed structures seen in the solubilized mutant constructs^12^. Gobeil et al.^11^ report that for the D614 S-protein construct with the furin site removed, the presence of the prolines produces structures, ACE2 binding, thermal stability, and antibody binding that are remarkably similar to the WT K986/V987 pair. In our equilibrium simulations this salt bridge has an ∼30% occupancy, averaged over all three protomers in the two independent trajectories. Reduction in the aforementioned salt bridge occupancy originates from competing with intra-protomer salt bridges between K986 and nearby acidic residues E748, E990, and D985, present in the ∼15-25% range, suggesting transient interaction between K986 and the RBD aspartic acids (D427/D428). These competing interactions are illustrated in Fig. S10.

We repeated the REUS simulation with the diproline mutation for the un-glycosylated system. The resulting PMF displays a modest energy difference between the down and up RBD states favoring the up state (Fig. S11), similar to that seen in the WT PMF (Fig. 2b) for these ancestral D614 constructs.

The similarity between the down-state and up-state energy wells compared with those for WT (Fig. 2b) suggests that the diproline mutation produces only a modest perturbation of their relative populations in these unglycosylated systems. In contrast to the minima, the barrier in the diproline mutant is seen to increase, suggesting a modulation of the opening kinetics. Given the transient nature of the K986 salt bridges and the similar relative energies of the down/up RBD states, an additional impact of the proline mutations is likely to be their tendency to break and distort α-helices. The HR1-CH turn region undergoes a large conformational change upon the transition from the preto post-fusion states, forming an extended α-helix^12^. The presence of the prolines, residues known to break/kink α-helices, inhibits the formation of the long α-helix characteristic of the post-fusion conformation^49^. Of note, recent simulations by Wang et al.^50^ exploring the transition from the preto post-fusion states have shown that while the WT sequence at 986/987 has a tendency to become helical, replacement with prolines disrupts the formation of this secondary structure.

### Glycans differentially impact epitope exposure

Currently, nearly ten thousand neutralising antibodies targeting the SARS-CoV-2 S-protein have been discovered. The most prominent target is the S-protein RBD^51,52^, although some target the NTD and other non-RBD epitopes^53^. According to the continuously updated CoV-AbDab database^54^, a total of 9906 (and growing) neutralizing antibodies target the S-protein, while only 3955 target non-RBD epitopes. One advantage of targeting non-RBD epitopes on the S-protein is that they can be recognized even when the RBD is in a down conformation^20,55^. In contrast, the RBD can only be identified in the up conformation owing to the strong glycan coverage of RBD in the down conformation^26^.

To understand the effect of spike-opening dynamics on epitope exposure, we calculated the accessible surface area of a number of epitopes along the MEP identified from REUS. We selected seven different antibodies^16,17,56–60^ with epitopes spanning multiple regions on RBD-A (including cryptic epitopes) and the NTD-B of the S-protein. Details of these antibodies are provided in the Supporting Information. We performed separate antibody accessible surface area (AbASA) calculations including and excluding the glycans using the same sampling approach with a 7-Å probe. A similar probe size^20^ as well as a smaller one^33^ were used in previous studies. While this moderate (7-Å) probe size may make some crevices appear slightly exposed^20^, it is optimal for cryptic epitopes^33^.

In general, the AbASA either remains the same or increases significantly when RBD-A transitions to the up state. The epitope of the STE90-C11^59^ antibody has a jump in AbASA during the closed-to-open transition. The CR3022 antibody binds to a cryptic epitope^57^, which is completely covered in the down state and is only slightly exposed in the up state. Protein residues cover this epitope similarly throughout the spike opening path for MEPs with and without glycans (Fig. S12). Remarkably, the AbASA of the 2-4 antibody does not change during the downto-up transition of the RBD, in spite of the epitope being located in the RBM (Fig. 6a). Therefore, we conclude that the entire RBM does not become exposed even in the up state. The AbASA for the epitope of the 4A8 antibody does not change significantly during spike opening since its epitope is located in the N-terminus^17^, which did not undergo conformational changes. The AbASA of S2M11’s epitope does not change along the spike opening path. Furthermore, part of the S2M11’s epitope is on the RBD of the adjacent protomer, which does not open in our simulations. The epitope of the S2M11 antibody^60^ slightly overlaps with the RBD of one protomer (residues 440, 441, and 444), while the majority coincides with the binding site of the ACE2 glycan at N322 on the RBD (residues 369, 371 to 374, and 440)^61^. Surprisingly, the epitope of the S309 antibody^58^ becomes less exposed in the open state without an increase in coverage by the glycans (Fig. 6b), indicating that a part of the RBD (Table S3) also become less exposed in the up state compared to the down state.

**Figure 6.**
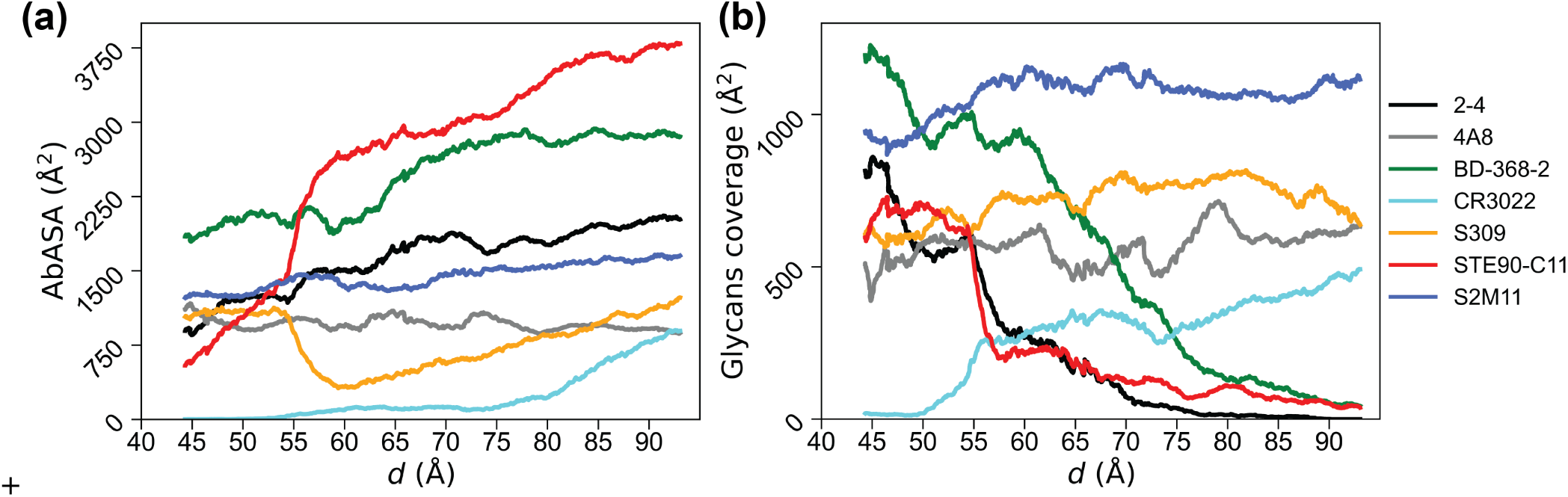
Epitope analysis for selected antibodies. (a) Exposed area on antibody epitopes (AbASA) in the presence of protein and glycans. (b) Surface area of epitopes covered by glycans along the MEP quantified by subtracting the two AbASA values calculated with and without glycans. All accessible surface area calculations were performed using a 7-Å probe.

Of note, the epitopes of antibodies chosen here are in the NTD and RBD of the spike, the primary regions of variant mutations with effective neutralization evasion^5,62,63^. Impacted antibodies include STE90-C11, 4-8, S2M11, BD-368-2 by Omicron BA.1^62,64,65^ and increased S309 escape by Omicron BA.2, BA.4/BA.5^63^. In the latter case, the loss of neutralization involves a region proximal to the N343 glycan, highlighting the important role of glycans in the immune response.

## Discussion

SARS-CoV-2 infection is initiated when an ACE2 receptor binds to the S-protein, which inter-converts between up and down states with only the up state allowing for ACE2 binding. Consequently, the energetics of the interconversion control the initiation of infection. To this end, we computed a two-dimensional energy landscape of the SARS-CoV-2 S-protein up-down inter-conversion. Using an aggregate total of 160 μs of REUS calculations, we elucidate the collective roles of S-protein glycans in its opening dynamics. Our results indicate that the free-energy barrier of spike opening is ∼7 kcal/mol higher in the presence of glycans compared to the fully un-glycosylated spike. While the free energy greatly favors the down state in the presence of the glycans, the energy difference between the two states of the S-protein diminished to only ∼0.5 kcal/mol without glycans. One may assume more up-state S-proteins would present a higher chance of binding ACE2 receptors and hence increase its viral fitness. However, there are a few opposing effects of having more up-state S-proteins as indicated by prior studies^26,66^ and reaffirmed by our analysis. The RBD in the down state is heavily shielded by glycans^20,26^, while in the up state, it is more exposed to neutralizing antibodies (Fig. 6a). Furthermore, ACE2 binding strongly depends on the binding affinity between ACE2 and the S-protein, which may involve interactions with glycans as well as protein. Therefore, a high up-state population does not necessarily reflect a high ACE2-bound-state population.

The atomic level description described here, and elsewhere^26–36^, becomes particularly relevant in light of the central role that dynamics plays in the functioning of the spike protein. Although cryo-EM structures have been a centerpiece in delineating these states, the transitions between them as well as intermediary states are unavailable.

Molecular dynamics provides a complementary tool to these experimental studies to achieve a more complete picture of the functioning of these proteins, including the kinetic effects of altered glycans. Moreover, even the relative populations of the down and up states are not without some uncertainty. For example, although the recent cryo-EM structure of the Omicron ectodomain variant has predominantly 1-up RBD^67^, alternate populations are reported^68^ and the full length spike appears to support a ensemble of 2-up/1-down RBDs^69^. Such diversity of conformations is likely a result of alternate constructs and/or experimental conditions^68^.

The epitopes of antibodies on the spike can be protected either by protein residues or by glycans, and the coverage provided by each changes in an epitope-dependent manner along the spike opening path. The STE90-C11 antibody has the greatest epitope exposure when the RBD is open, while the BD-368-2 antibody has an epitope that is always significantly exposed, irrespective of the RBD’s position. Additionally, the exposure of the BD-368-2 epitope is very similar to the epitope of STE90-C11 in the up state. Therefore, these two antibodies show very high efficiency in neutralizing SARS-CoV-2 S-protein, both with sub-nanomolar IC_50_ values^56,59^. The CR3022 antibody binds to a cryptic epitope; consequently, its AbASA is smaller compared to other epitopes. However, the lack of exposed surface area, even in the up state, is compensated for by strong electrostatic interactions at the epitope-CR3022 interface^70^. Note that the S309 Fab binds to a proteoglycan epitope, which includes the glycan at N343. However, since most of the S309 epitope is not part of the RBM, it may attach to the spike protein in both down and up states^58^. However, the effect may be nullified through mutations near the N343, as observed by the Omicron escape^63^.

We analyzed interactions of individual glycans with different domains of the S-protein using the two MEPs obtained, one with and one without glycans. The glycans at N122 and N165 disrupt the hydrogen bonds between the opening RBD-A and the neighboring NTD-B, destabilizing the up state and hence pushing the equilibrium population towards the down state when compared to the un-glycosylated system. The glycans at N165 and N343 also stabilize the RBD-A down state by wrapping around the RBM. Conversely, the glycan at N343 serves as the so-called “glycan gate” propping up the RBM^32^, while the glycan at N165 helps by supporting the up-state RBD-A with the aid of the glycan at N343. Consequently, these two glycans stabilize both down and up states, creating a local minimum for each. The complicated role of the glycan at N165 may explain conflicting experimental results regarding whether the N165A mutation increases or decreases spike binding^26,71^.

The spike protein is the primary target of neutralizing antibodies, presenting an assortment of epitopes in the down and up states, and is the basis for current vaccines and boosters^5,15–17^. Under evolutionary pressure the spike is rapidly mutating, with the emergence of more transmissible variants with increased antibody escape^5^. Although the virus continues to mutate, to date, vaccines and boosters appear to provide adequate protection. An effective approach to treat the prolonged pandemic continues to be identification and evaluation of important immunological regions on the spike, both in the down and up states as well as transitions between them, particularly as it appears the spike will continue to mutate and potentially compromise the current vaccines/boosters. Given the altered structure and dynamics reported for SARS-CoV-2 spike variants^11,25,67,68^, a detailed description of the energetics and kinetics for the opening of the spike protein is necessary, and computational approaches as described here provide a valuable complementary tool in the development of effective treatments.

In closing, we calculated the energetics along the spike-opening path for SARS-CoV-2 S-protein with atomic-level insight into the roles of glycans in the opening process. Furthermore, we highlighted how the spike-opening energetics impacts the kinetics of ACE2 binding and epitope exposure. These findings, especially the conformations along the spike-opening path, will facilitate the design of effective nanobodies and antibodies to fight the ongoing COVID-19 crisis.

## Materials and Methods

### Model building

We modeled the up and down states based on the cryo-electron microscopy (cryo-EM) structures by Walls et al.^10^ (PDB: 6VYB) and by Cai et al.^12^ (PDB: 6XR8). The down state structure reported by Cai et al. is a detergent purified full-length WT S-protein construct at 2.9 Å resolution^12^. It exhibits several critical differences with earlier reported structures: (i) more resolved NTD with an additional glycosylation site at N17, (ii) the WT sequence at the central helix/loop region, rather than the pre-fusion stabilizing 2PP mutation, and (iii) an approximately 25-residues-long segment, residues 828-853, that were previously unresolved. This latter region is adjacent to a critical lysine (K854) that provides a salt bridge partner for D614. The loss of this salt bridge has been implicated in the increased infectivity of the D614G SARS-CoV-2 strains. Furthermore, Cai et al.^12^ also resolved two disulfide bonds that were previously missing, one in the N-terminal (C15-C136) and another in the central helix/loop region (C840-C851). In our model, the missing residues in the down-state structures were added with SWISS model^72^ using the 6XR8 structure as a template. The model was cleaved at the furin cleavage site between residues R685 and S686. The S1 part, which contains the RBD and NTD, is more closely packed in the 6XR8 structure compared to the 6VXX, and the S2 part, which contains the HR1 and CH, is aligned between them^12^. The S1 and S2 part of our model include residues N14-R685 and S686-S1147, respectively. Ten disulfide linkages are added in the S1 part between residues C15-C136, C131-C166, C291-C301, C336-C361, C379-C432, C391-C525, C480-C488, C538-C590, C617-C649, and C662-C671 and five are added to the S2 part C738-C760, C743-C749, C840-C851, C1032-C1043, and C1082-C1126. The missing parts in the up state were modeled using the minimized down-state model. Glycosylation sites are located on N17, N61, N74, N122, N149, N165, N234, N282, T323, N331, N343, N603, N616, N657, N709, N717, N801, N1074, N1098, and N1134. Overall, there are 19 N-linked and 1 O-linked glycans present in each protomer resulting in a total of 60 glycans for one S-protein trimer model. The glycan at each site with the highest population in the mass spectroscopy data by Crispin and coworkers^19^ was added to the site using the GLYCAM Web server developed by the Woods group (http://glycam.org)^37,38^. The glycan compositions are illustrated in Fig. S13. Missing hydrogen atoms were added to all systems, after which they were solvated in a 195 × 193 × 212 Å^3^ water box. We added Na^+^ and Cl^*−*^ ions to achieve a salt concentration of ∼0.150 M. The number of atoms in protein, water, glycan, and ions were kept same for REUS calculations. The systems (up or down states) contain a total of 758,531 atoms (63,312 protein and glycan atoms, 661 Na^+^, 652 Cl^*−*^) and 633,864 atoms (52,476 protein atoms, 549 Na^+^, 546 Cl^*−*^) with and without glycans, respectively.

### Molecular dynamics (MD) simulations

All simulations were performed using NAMD 2.14^73,74^ with the CHARMM36m protein force field^75^, CHARMM36 glycan force field^76^, and TIP3P water^77^. Each system was equilibrated for 5 ns with the temperature and pressure fixed at 310 K and 1 atm, respectively, using Langevin dynamics and piston^78^, respectively. A uniform 4-fs time step was employed through the use of hydrogen mass repartitioning (HMR)^79,80^. The long-range electrostatics were calculated every time step using the particle-mesh Ewald method^81^. A short-range cutoff for Lennard-Jones interactions was set at 12 Å, with a switching function beginning at 10 Å. Bonds involving hydrogen atoms were constrained to their equilibrium length, employing the SETTLE algorithm for water molecules and the SHAKE algorithm for all others.

We also ran metadynamics, simulated annealing and REUS simulations. Additional details are provided in the Supporting Information.

## Supporting information

Spike opening path

Close-up of glycans around RBD during opening

## Acknowledgements

An award of computer time on the Summit supercomputer at Oak Ridge National Laboratory was provided through the COVID-19 High Performance Computing Consortium. Additional computational resources were provided through the Extreme Science and Engineering Discovery Environment (XSEDE; TG-MCB130173), which is supported by the National Science Foundation (NSF; ACI-1548562). This work also used the Hive cluster, which is supported by the NSF under grant number 1828187 and is managed by the Partnership for an Advanced Computing Environment (PACE) at the Georgia Institute of Technology. The authors thank Mahmoud Moradi for his guidance in calculating the mean first passage time. Y.T.P. is supported by a Texas Advanced Computing Center (TACC) Frontera Fellowship. Frontera is supported by NSF grant No. OAC-1818253.

## Author contributions

Y.T.P., A.A., D.L.L., and J.C.G. designed research; Y.T.P., A.A., D.L.L., and A.P. performed research and analyzed data; and all authors wrote the manuscript.

## Supplementary Information

Details of modeling steps, glycosylation, collective variables, definitions of epitopes, additional analysis, and two movies are provided in the SI.

## Ethics declarations

Competing interests: The authors declare no competing interests.

## Data availability

S-protein conformations along the MEP for each of the three PMFs are available at the NSF MolSSI COVID-19 Molecular Structure and Therapeutics Hub at https://covid.molssi.org

