## Supporting Information for "SARS-CoV-2 spike opening dynamics and energetics reveal the individual roles of glycans and their collective impact"

#### Details of molecular dynamics (MD) simulations

**Metadynamics simulations.** After equilibration, we ran two independent metadynamics simulations along the two collective variables $d$ and $\phi$ (defined in Main Text Fig. 1) for each system (WT glycosylated, WT un-glycosylated and un-glycosylated with diproline mutation), one starting from down state and one from the up state. The colvars module of NAMD was used to construct all collective variables[1]. The biases were deposited with a hill-height of 0.2 kcal/mol, a width of 1.0 Å and 1.0° for $d$ and $\phi$, respectively, and a rate of 1 ps$^{-1}$. The systems were confined to the $d$ and $\phi$ within the range of the cryo-EM structures of the down (PDB: 6XR8) and up (PDB: 6VYB) states. The temperature was held at 310 K using Langevin dynamics and the volume as fixed. A uniform 4-fs time step was employed through the use of hydrogen mass repartitioning (HMR)[2,3]. Other simulation parameters were identical to those used in the equilibrium simulations.

**REUS simulation seeding.** We then used the metadynamics trajectories to seed the REUS runs according to the following procedure. For each system, two preliminary PMFs were generated, one from the down-state metadynamics and another from the up-state one. The conformational space was broken into small grids and each grid became a single REUS window in the next stage. Windows that have their PMFs from both the down- and up-state metadynamics below a certain threshold (Table S2) were dropped and would not be used in the REUS simulation. For the remaining grids, we picked a snapshot from either the down- ($F_{\text{down}}$) or up-state ($F_{\text{up}}$) metadynamics trajectory according to the Boltzmann weight of their respective PMF.

$$P_{\text{down/up}}(d,\phi) = \frac{\exp\left[-\beta F_{\text{down/up}}(d,\phi)\right]}{\exp[-\beta F_{\text{down}}(d,\phi)] + \exp\left[-\beta F_{\text{up}}(d,\phi)\right]} \tag{S1}$$

If $P_{\text{down}} > P_{\text{up}}$, a snapshot from the down-state metadynamics trajectory was picked for that grid and vice versa. With this, 387 windows (181 from down, 206 from up) were selected for the WT glycosylated system; 392 windows (79 from down, 313 from up) for the WT un-glycosylated system; and 353 windows (119 from down, 234 from up) for the un-glycosylated system with diproline mutation.

**Simulated annealing of glycans.** For the glycosylated system, we further performed simulated annealing on the glycans for all snapshots extracted from the metadynamics simulations. Starting at 1000 K, the systems were slowly cooled down to physiological temperature at steps of 823 K, 677 K, 557 K, 458 K, 377 K, and 310 K. Each temperature was run for 2 ns. During the simulation, all protein atoms were fixed and the glycans, water molecules and ions were allowed to move freely. Due to the high velocities of the atoms at 1000 K and 823 K, a time step of 2 fs was used together with HMR. For temperatures lower than or equal to 677 K, a time step of 4 fs was used with HMR. Other simulation parameters were identical to those used previously. The final ring conformations of the glycans were confirmed to be consistent with the expected values (Fig. S14).

**REUS simulations.** With the initial configurations prepared, we ran REUS simulations for each system along $d$ and $\phi$ within the regions selected according to the metadynamics simulation PMF. The ranges of $d$ and $\phi$ are from 44.0 Å to 72.5 Å and –56.0° to 1.0°, respectively. The restraining force constants along $d$ and the window centers were adjusted to ensure sufficient overlap and exchange between neighboring windows. The force constants along $\phi$ were kept constant at 1.0 kcal $mol^{-1}deg^{-2}$. The REUS simulations of the WT glycosylated system, WT un-glycosylated system and un-glycosylated system with diproline mutation were run for 32 ns, 36 ns and 29 ns, respectively. Simulation parameters were identical to those used in the metadynamics simulations. The sampling data from REUS was used to calculate the PMF for each system using Multistate Bennett Acceptance Ratio (MBAR), implemented in the Python module pymbar[4]. For the WT glycosylated and un-glycosylated systems, we ran three more rounds of REUS simulations using snapshots extracted from the previous rounds in a similar manner as for the metadynamics simulations. For the second round, the simulation regions were re-selected using a new energy cutoff based on the PMFs computed from the first round of REUS (Table S2). The restraining force constants and the windows centers were also recalibrated. The second round of REUS for the WT glycosylated and un-glycosylated systems were run for 37 ns and 56 ns, respectively. In light of the result from the second round of REUS, we ran a third round of REUS, expanding the PMF region to fully cover the two energy wells. Two independent REUS simulations were run during the third round for each of the energy wells. The ranges of $d$ and $\phi$ were from 40.5 Å to 51.0 Å and from –66.5° to –21.5°, respectively, for the down-state REUS. The ranges of $d$ and $\phi$ were from 57.5 Å to 99.5 Å and from –35.5° to 45.5°, respectively, for the up-state REUS. New metadynamics simulations were run to seed regions never sampled by previous REUS simulations. Again, windows were selected based on a new energy cutoff (Table S2), and the restraining force constants and the window centers were calibrated to maximize overlap and exchanges between neighboring windows. The third rounds of REUS for the WT glycosylated down- and up-state systems were run for 16 ns and 12 ns, respectively, and those for the un-glycosylated system were 33 ns and 28 ns, respectively. Finally, a last round of REUS was run to cover the full range from down to up state. All the windows from rounds 2 and 3 were adopted, with a few added on the edges to make sure all low-energy regions were covered (Table S2). The REUS for the WT un-glycosylated system was run for 36 ns directly using the last frames from previous rounds of REUS. For the glycosylated system, we generated new seeding structures from the un-glycosylated REUS for windows around the energy barrier to help bridge the gap between the down- and up-state structures in the glycosylated REUS simulation. Using the structures from previous rounds of REUS and the newly generated structures from the completed un-glycosylated REUS simulation, the glycosylated REUS was run for 36 ns. The data presented in the Main Text for the WT systems are based on the combination of data from the second to the last round of REUS. The total aggregation time of simulation data used for the WT glycosylated system, WT un-glycosylated system and un-glycosylated system with diproline mutation REUS were 65 µs, 91 µs, and 12 µs, respectively.

### Analysis

**Minimum energy path (MEP).** The MEPs were computed using the algorithm by Ensing et al.[5] and the code implemented by Mahmoud Moradi[6]. The two energy minima and the saddle point (the point with lowest energy to cross an energy barrier) were found using a separate in-house python script, and the MEP from the saddle point to each energy minimum was found using the code by Moradi. The paths were smoothed by the least-square fitting B-spline function from python library SciPy[7].

**Mean first passage time (MFPT).** Let $\lambda(\mathbf{d}, \theta)$ be the one-dimensional path parameter that describes the position of the system along the MEP. Assuming the dynamics along $\lambda$ can be effectively described by a diffusive model, we may apply the Smoluchowski diffusion equation to describe the process[8,9]:

$$\frac{\partial}{\partial t}p(\lambda,t|\lambda_0,0) = \frac{\partial}{\partial \lambda}D(\lambda)e^{-\beta F(\lambda)}\frac{\partial}{\partial \lambda}\left(e^{\beta F(\lambda)}p(\lambda,t|\lambda_0,0)\right) \tag{S2}$$

where $p(\lambda,t|\lambda_0,0)$ is the probability of finding the system at $\lambda$ after time $t$, given it was at $\lambda_0$ at time 0, $D(\lambda)$ is a position-dependent diffusion constant, and $F(\lambda)$ is the free energy at $\lambda$. By rearranging the terms, the MFPT $\bar{\tau}_{FP}$, or the rate inverse $k^{-1}$, from the initial (A) to the final (B) state is given by:

$$k^{-1} = \bar{\tau}_{FP} = \int_{\lambda_A}^{\lambda_B}\frac{1}{D(\lambda)}e^{\beta F(\lambda)}\int_{\lambda_0}^{\lambda}e^{-\beta F(\lambda')}\,\mathrm{d}\lambda'\,\mathrm{d}\lambda \tag{S3}$$

While $F(\lambda)$ was readily available from the PMF obtained through REUS, $D(\lambda)$ was approximated using a generalized-Langevin-equation-based method, derived by Roux and co-workers and implemented by Gaalswyk et al.[10]. From a time series of a coordinate $x_k$ with the system simulated under a harmonic restraint, the method computes the diffusion constant $D_k$ along $x_k$ by relating it to its velocity autocorrelation function (VACF). With a series of coordinate transformations[11], $D_k$ is transformed from the Cartesian space to the collective variable space ($D_{ij}$), and then to the path variable space ($D(\lambda)$), which was then inserted back into Eq. S3.

$$D_{ij} = \sum_k D_k\left\langle\frac{\partial z_i}{\partial x_k}\frac{\partial z_j}{\partial x_k}\right\rangle, \quad D(\lambda) = \sum_{ij}D_{ij}\frac{\partial \lambda}{\partial z_i}\frac{\partial \lambda}{\partial z_j} \tag{S4}$$

To determine $D_k$ for each atom and each of its coordinates involved in the definition of the collective variables $d$ and $\phi$, we ran a 1-ns simulation for each window along the MEP, with all $C_\alpha$ atoms of the protein restrained by a force constant of 5 kcal mol$^{-1}$Å$^{-2}$. A 2-fs time step was used without the application of HMR. Other simulation parameters were identical to the REUS simulations.

**Kinetics analysis.** We modeled the down-to-up transition and subsequent binding of the RBD to ACE2 according to the chemical equation

$$\mathrm{D} + \mathrm{ACE2} \underset{k_\mathrm{close}}{\overset{k_\mathrm{open}}{\rightleftharpoons}} \mathrm{U} + \mathrm{ACE2} \underset{k_\mathrm{off}}{\overset{k_\mathrm{on}}{\rightleftharpoons}} \mathrm{U{:}ACE2}$$

The associated master equation is

$$\frac{\mathrm{d[D]}}{\mathrm{d}t} = -k_\mathrm{open}[\mathrm{D}] + k_\mathrm{close}[\mathrm{U}] \tag{S5}$$

$$\frac{\mathrm{d[ACE2]}}{\mathrm{d}t} = k_\mathrm{off}[\mathrm{U{:}ACE2}] - k_\mathrm{on}[\mathrm{U}][\mathrm{ACE2}] \tag{S6}$$

$$\frac{\mathrm{d[U]}}{\mathrm{d}t} = k_\mathrm{open}[\mathrm{D}] - k_\mathrm{close}[\mathrm{U}] - k_\mathrm{on}[\mathrm{U}][\mathrm{ACE2}] + k_\mathrm{off}[\mathrm{U{:}ACE2}] \tag{S7}$$

$$\frac{\mathrm{d[U{:}ACE2]}}{\mathrm{d}t} = -k_\mathrm{off}[\mathrm{U{:}ACE2}] + k_\mathrm{on}[\mathrm{U}][\mathrm{ACE2}] \tag{S8}$$

The rates $k_\mathrm{open}$ and $k_\mathrm{close}$ come from our own calculations. For the glycosylated spike, $k_\mathrm{open} = 0.68$/s and $k_\mathrm{close} = 1.17 \times 10^3$/s, while for the un-glycosylated spike, $k_\mathrm{open} = 7.01 \times 10^3$/s and $k_\mathrm{close} = 2.01 \times 10^3$/s. The rates $k_\mathrm{on} = 1.40 \times 10^6$/M·s and $k_\mathrm{off} = 6.54 \times 10^{-3}$/s were taken from Lan et al.[12] for the RBD alone in order to isolate the effects of binding from conformational changes in the spike. The system of first-order ordinary differential equations was numerically solved up to 250 s, for which all systems reached steady-state populations, using Mathematica along with the initial conditions $[\mathrm{D}]_i$ = 1 nM, $[\mathrm{U}]_i$ = 0 nM, $[\mathrm{U{:}ACE2}]_i$ = 0 nM, and $[\mathrm{ACE2}]_i$ = 15 nM. The concentration chosen for ACE2 is the same order of magnitude as $K_\mathrm{D} = 4.67$ nM, and we note that the qualitative results did not change for different values of $[\mathrm{ACE2}]_i$ although absolute populations did shift.

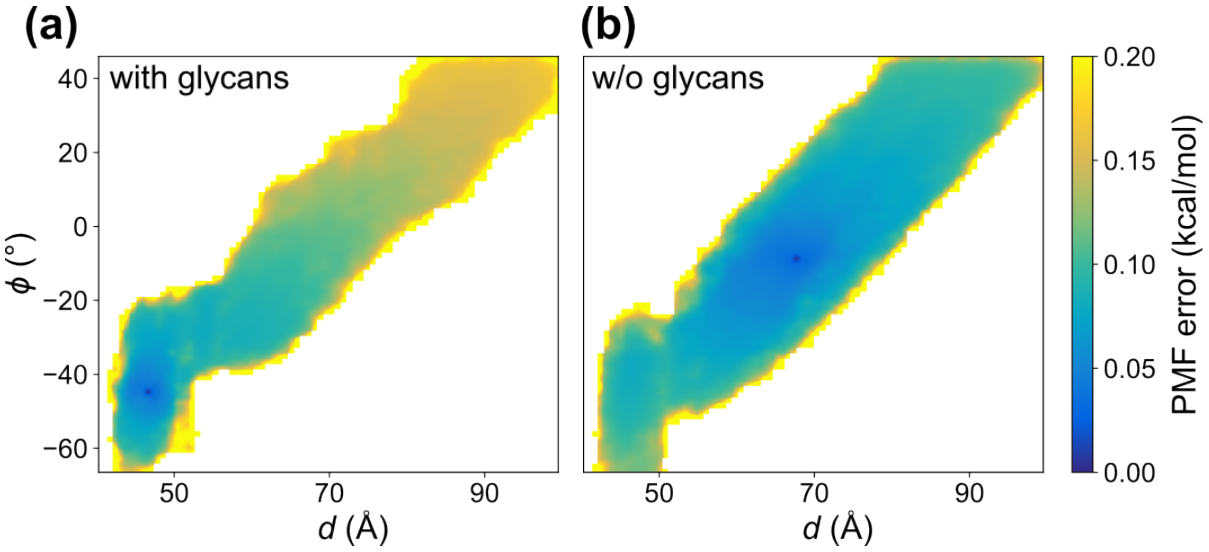


**Figure S1.** Uncertainties of the free energy differences with the lowest point on the PMF computed by MBAR for the (a) glycosylated and (b) un-glycosylated system.

**Hydrogen bond analysis.** The number of hydrogen bonds formed between RBD-A and other domains of the spike in the REUS trajectories were measured using the HBonds Plugin of VMD. When the distance between a hydrogen-bond donor atom (D) and an acceptor atom (A) is below 3.5 Å and the angle D-H-A is less than 35° from 180°, a hydrogen bond was considered to be formed between D and A. The number of hydrogen bonds was averaged over all frames from the same REUS window.

**Contact analysis.** Contact calculations were performed using a cutoff distance of 3.5 Å. In other words, when two heavy atoms from two different selections come within 3.5 Å, we count that as one contact. For the analysis done in Figs. S4 and S8, the number of contacts was averaged over all frames from the same REUS window. For the analysis done in Fig. S8, we separated conformations from the MEP in two categories and the number of contacts were averaged separately for each category. The two categories were defined by the $d$ values. The conformations with $d \leq 54.9$ Å were considered as the down state, while the rest were taken as the up state.

**Antibody accessible surface area (AbASA).** AbASA calculations were performed using the solvent-accessible surface area measurement tool as implemented in VMD with a 7-Å probe for each frame along the MEP. For the glycosylated system, we performed two accessible surface area calculations for each frame around each epitope using the *-restrict* option: AbASA of (calculation #1) protein and (calculation #2) protein + glycans. The difference in the accessible surface area obtained from these two calculations gives the coverage provided by the glycans only.

**Supplementary Videos.** One video shows the MEP for the spike with (left) and without (right) glycans. The other video illustrates the motion of key glycans along the MEP with the glycans rendered red (N122), green (N165), yellow (N234), and orange (N343).

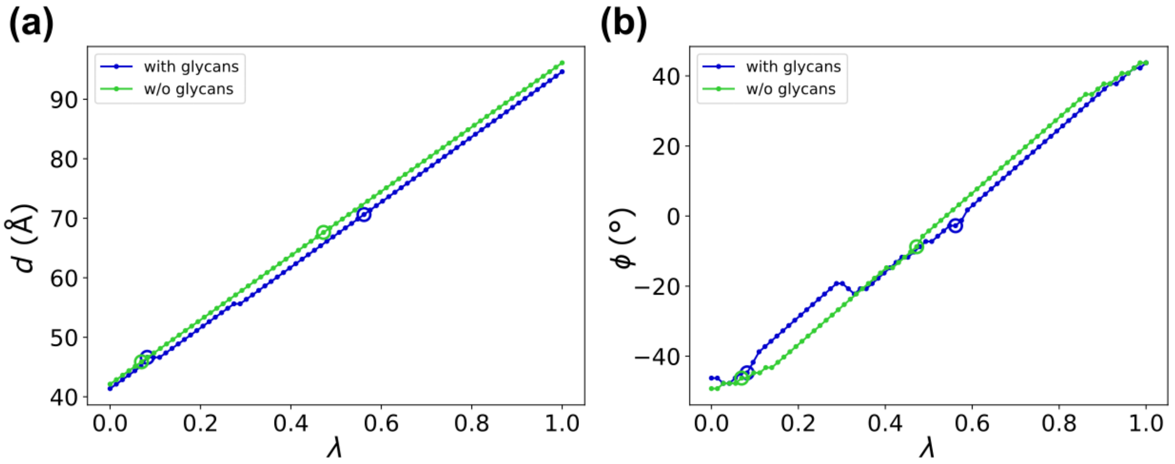


**Figure S2.** Path parameter $\lambda$. (a,b) The path parameter is defined as a value going from zero to one along the MEP from the bottom-left to the top-right corner of the PMF. The corresponding $d$ and $\phi$ are plotted against $\lambda$ in (a) and (b), respectively. The position of the down- and up-state energy minima are marked as empty circles.

| | glycosylated | | | un-glycosylated | | |
|---|---|---|---|---|---|---|
| Domains | x (Å) | y (Å) | z (Å) | x (Å) | y (Å) | z (Å) |
| SD1-B | $-31.6 \pm 1.1$ | $14.1 \pm 1.1$ | $-13.1 \pm 1.0$ | $-31.4 \pm 0.8$ | $14.7 \pm 1.2$ | $-12.4 \pm 1.0$ |
| SD1-A | $2.2 \pm 1.4$ | $-34.2 \pm 1.1$ | $-11.8 \pm 1.6$ | $2.6 \pm 1.7$ | $-34.8 \pm 1.2$ | $-10.9 \pm 2.1$ |
| SD2-A | $26.5 \pm 1.4$ | $-25.4 \pm 1.6$ | $11.6 \pm 1.2$ | $25.6 \pm 1.2$ | $-25.3 \pm 1.5$ | $12.5 \pm 1.0$ |
| NTD-A | $45.8 \pm 1.7$ | $-11.0 \pm 1.5$ | $-21.8 \pm 0.9$ | $45.2 \pm 1.6$ | $-11.2 \pm 1.6$ | $-21.7 \pm 1.1$ |

**Table S1.** The average position of the stationary domains used to define the two collective variables throughout the REUS simulations and their standard deviations.

| | PMF cutoff (kcal/mol) | | | | | Number of windows | | | | |
|---|---|---|---|---|---|---|---|---|---|---|
| Round | 1 | 2 | 3 | | 4 | 1 | 2 | 3 | | 4 |
| | | | Down | Up | | | | Down | Up | |
| WT with glycans | 30 | 28 | 25 | 26 | $\infty$ | 387 | 422 | 178 | 700 | 1049 |
| WT w/o glycans | 24 | 12 | 25 | 25 | $\infty$ | 392 | 356 | 257 | 669 | 1211 |
| diproline mutant w/o glycans | 28 | – | – | – | – | 353 | – | – | – | – |

**Table S2.** The PMF cutoffs for a window to be retained for the next stage of REUS simulation and the resulting number of REUS windows.

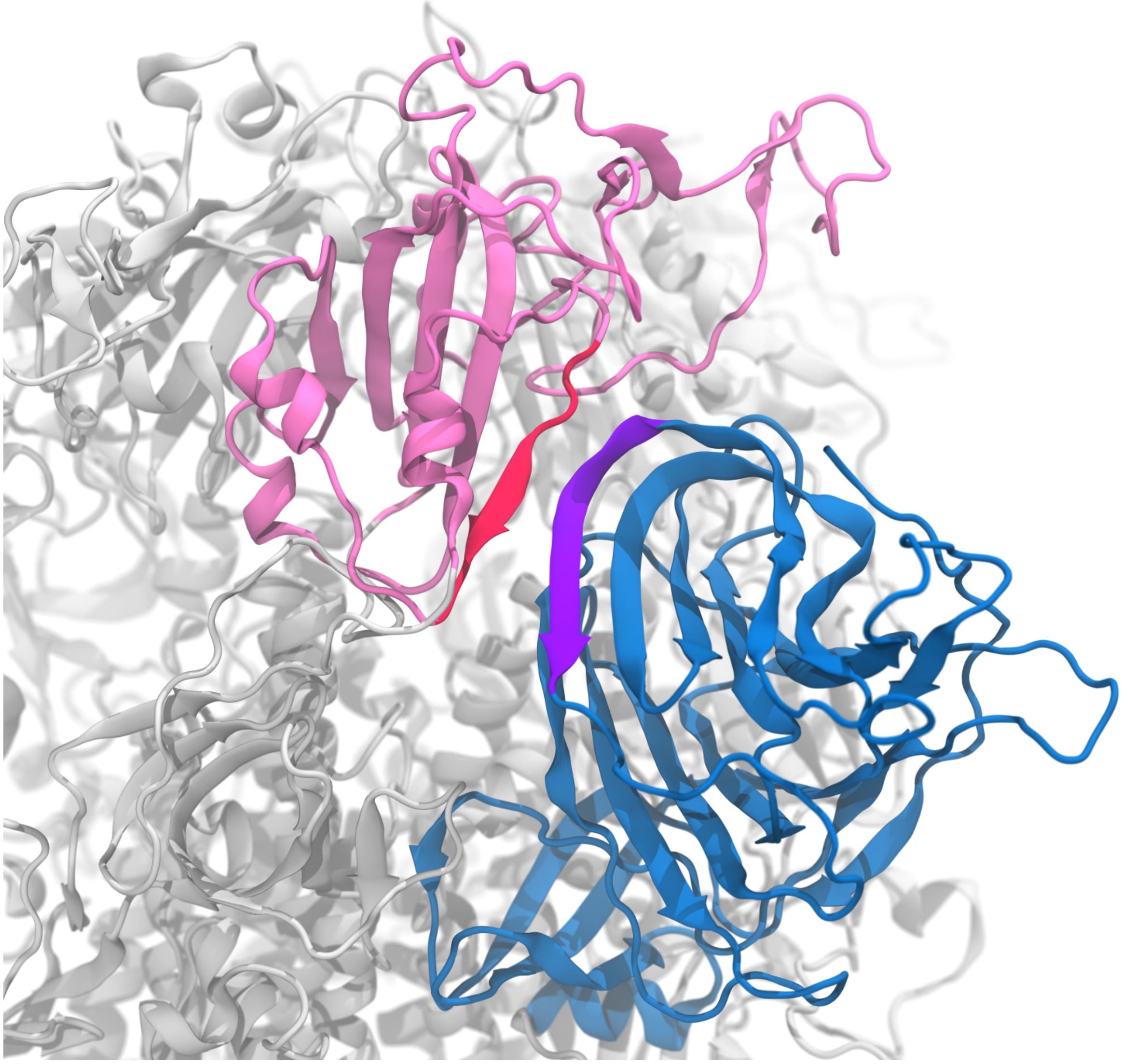

**Figure S3.** Snapshot from the un-glycosylated REUS simulation showing the β-sheets of RBD-A (pink) and NTD-B (blue) align with each other and form strong hydrogen bonds when the RBD-A is in the up state.

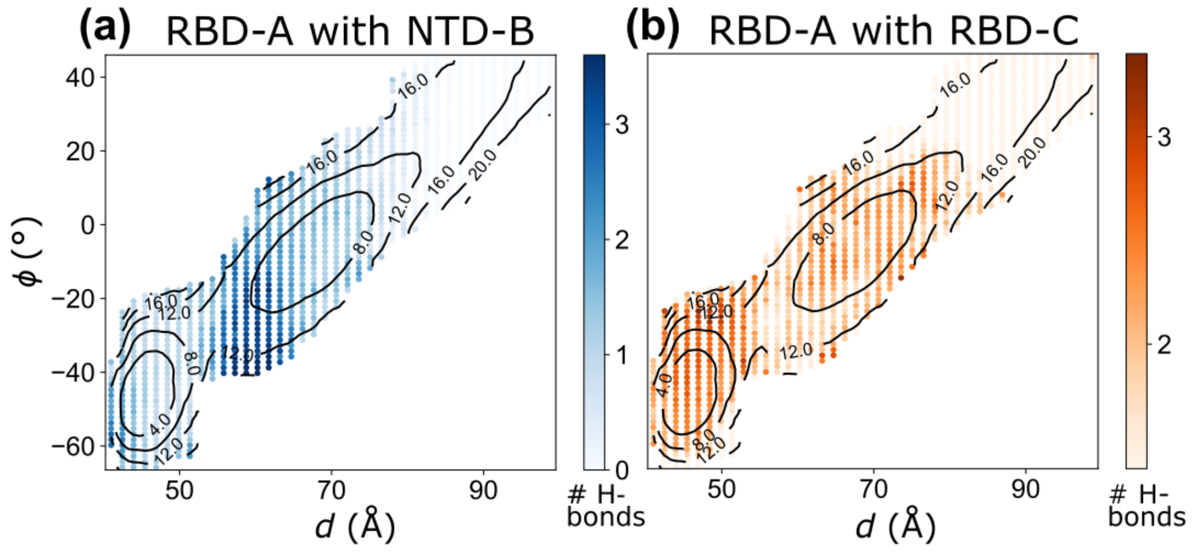


**Figure S4.** Hydrogen bond analysis for the glycosylated system. The average number of hydrogen bonds formed (a) between RBD-A and NTD-B and (b) between RBD-A and RBD-C are plotted against the two collective variables, $d$ and $\phi$. Contour lines of the PMF of the glycosylated system are plotted on top to show the location of the energy barrier.

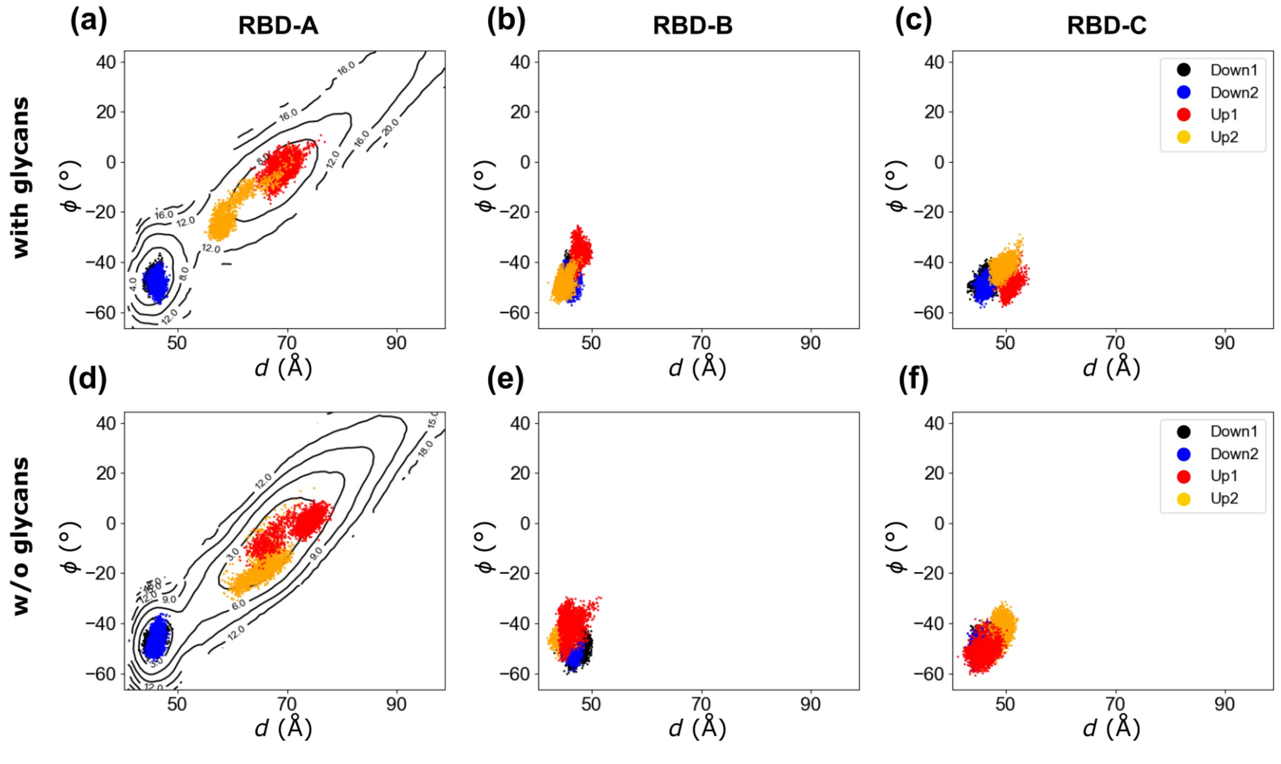


**Figure S5.** Collective variable $d$ vs. $\phi$ for the RBDs in $2 \times 2$-μs equilibrium simulations. (a-c) Glycosylated system. (d-f) Un-glycosylated system. The S-protein protomers in the down state, including (a,d) RBD-A that started at the down state (black/blue), (b,e) RBD-B, and (c,f) RBD-C, all remained in the free energy well of the down state for all replicas, with or without glycans. The up-state RBD-A (red/yellow) was less stable in comparison.

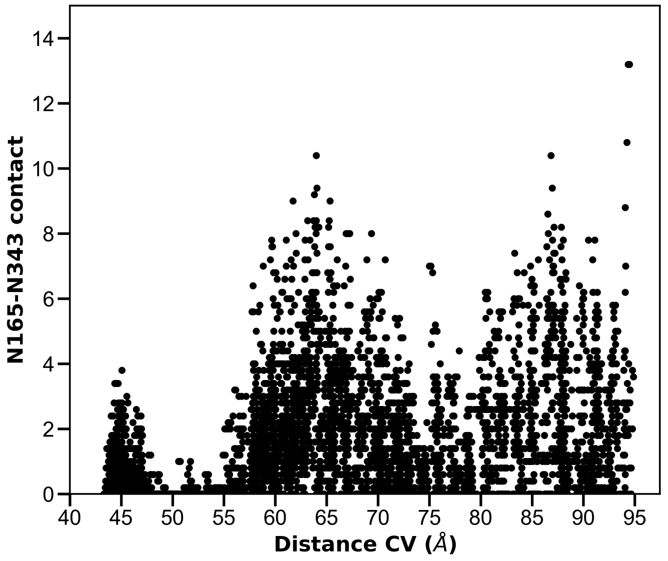


**Figure S6.** Contact between glycans at N165 and N343 along the MEP.

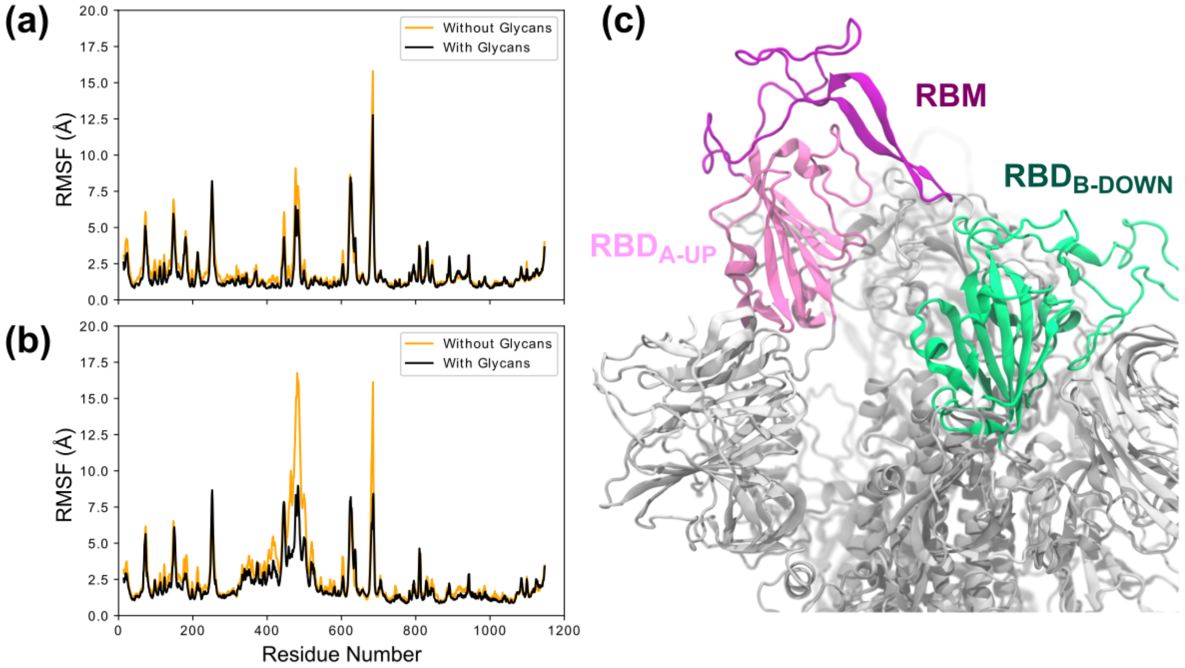


**Figure S7.** Root-mean-square fluctuation (RMSF) for equilibrium simulations. (a) RMSF of the down protomers, averaged over the 3 protomers and 2 replicas. (b) RMSF of the single up protomer, averaged over both replicas. RMSF computed after alignment and averaging over all $C_\alpha$ atoms of the trimer. (c) Snapshot from the equilibrium simulation without glycans, showing the RBM region (438–506) of the up-state protomer attached to the neighboring RBD-B.

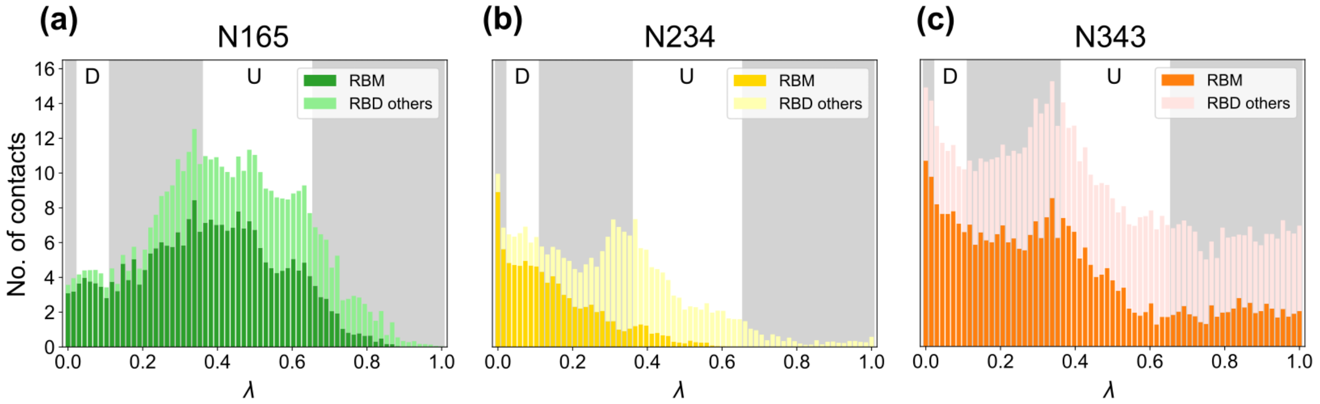


**Figure S8.** The number of contacts along the MEP between the glycans at (a) N165, (b) N234 and (c) N343, and RBD-A, which is separated into the RBM (439–506) and the non-RBM.

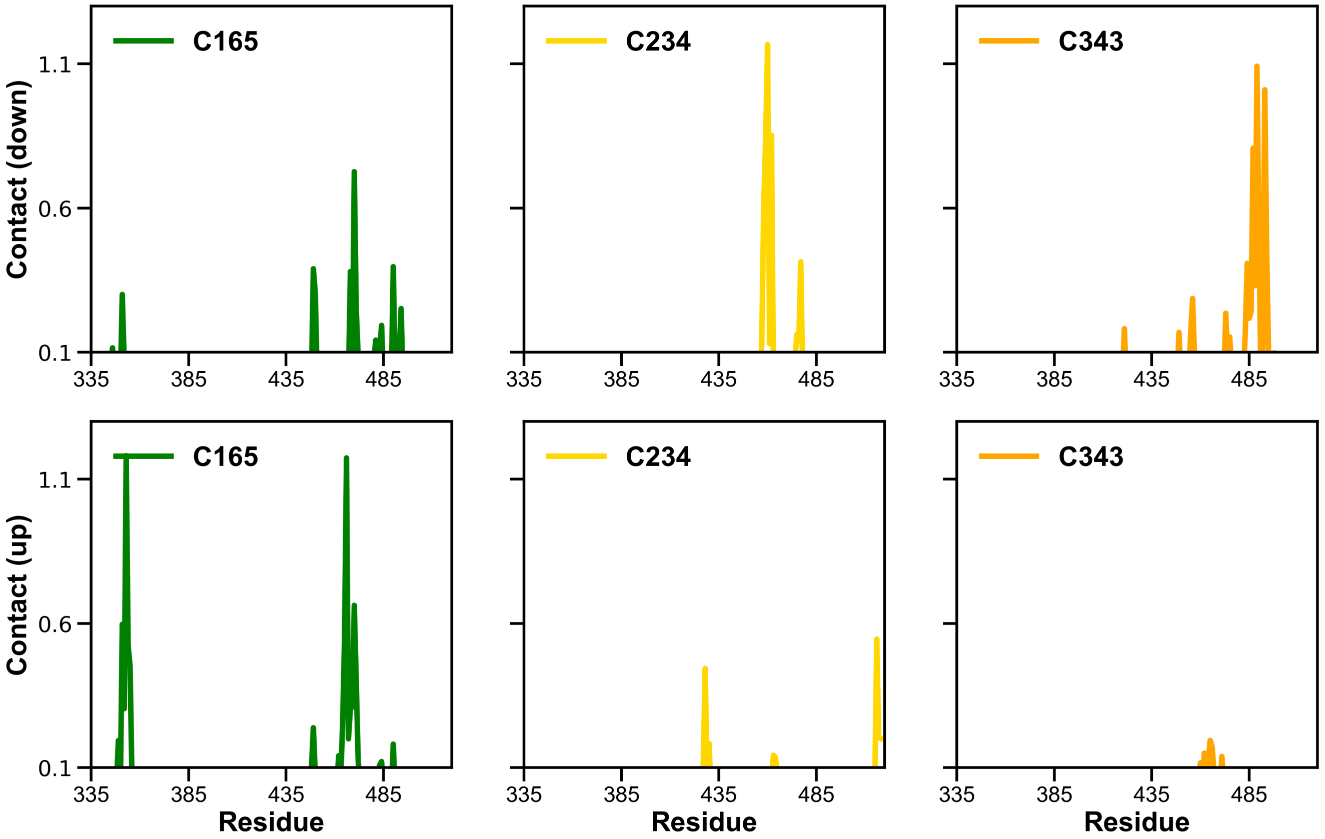


**Figure S9.** Average contact between glycans and RBD residues in up and down conformations of the MEP. Conformations in the range 43.1 Å $\leq d \leq$ 48.9 Å were considered as the down state, while the conformations with 60.1 Å $\leq d \leq$ 75.6 Å were taken as the up state.

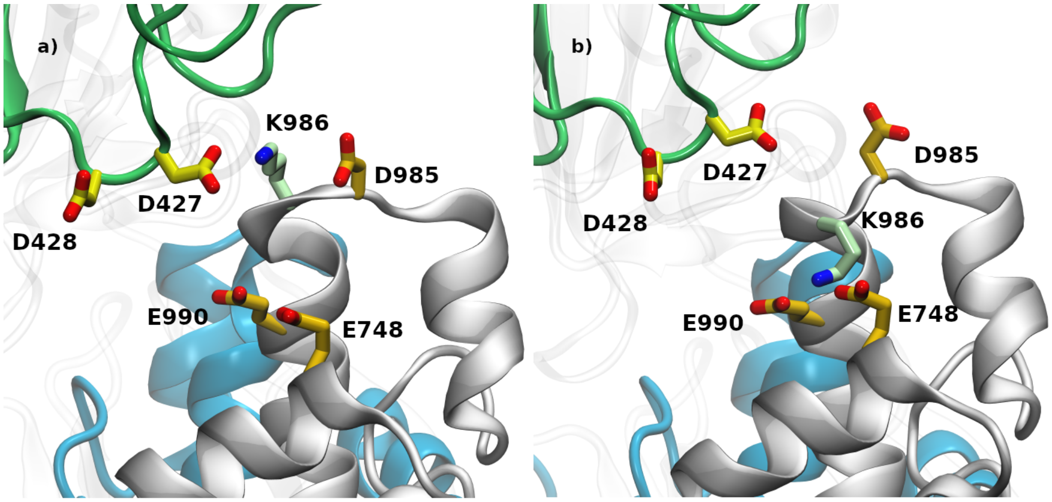


**Figure S10.** Intra-protomer and inter-protomer K986 interactions. K986 transiently interacts with a) D427 from a neighboring protomer or b) D985, E748, and E990 from the same one. D427, D428, and the green ribbon are from protomer B while K986, D985, E748, E990, as well as the silver and blue ribbons, are from protomer A.

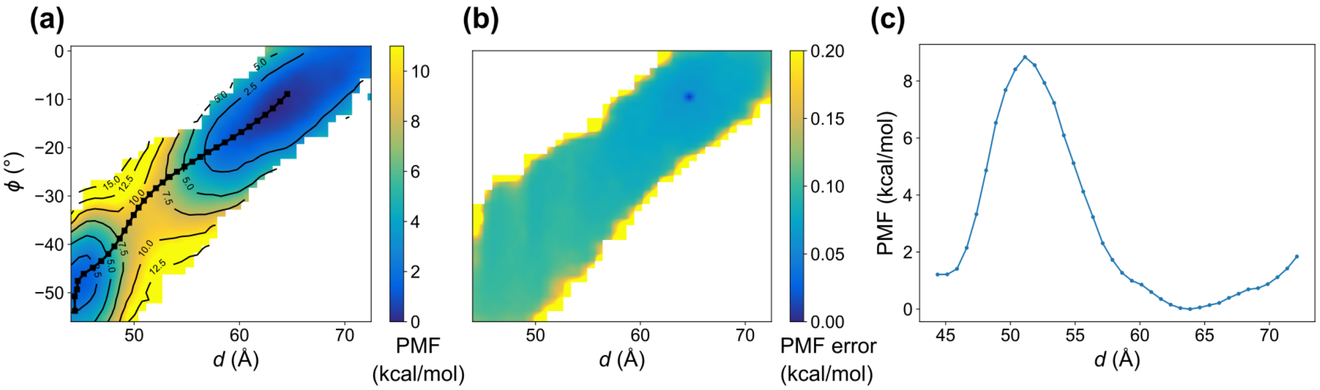


**Figure S11.** PMF of the diproline-mutated un-glycosylated spike. (a) The 2D PMF along two collective variables, $d$ and $\phi$. The black lines show the MEP. (b) Uncertainties of the free energy differences with the lowest point on the PMF computed by MBAR. (c) The free energies are projected onto $d$ and plotted as an 1D PMF.

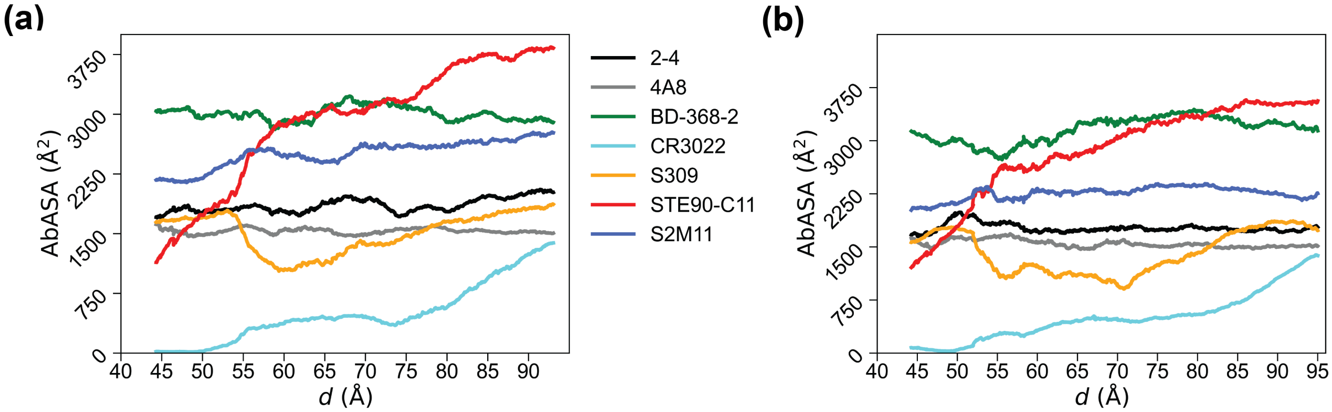


**Figure S12.** Epitope analysis of selected antibodies. Exposed area on antibody epitopes without glycans along the MEP obtained (a) with glycans and (b) without glycans. All accessible surface area calculations were performed using a 7-Å probe.

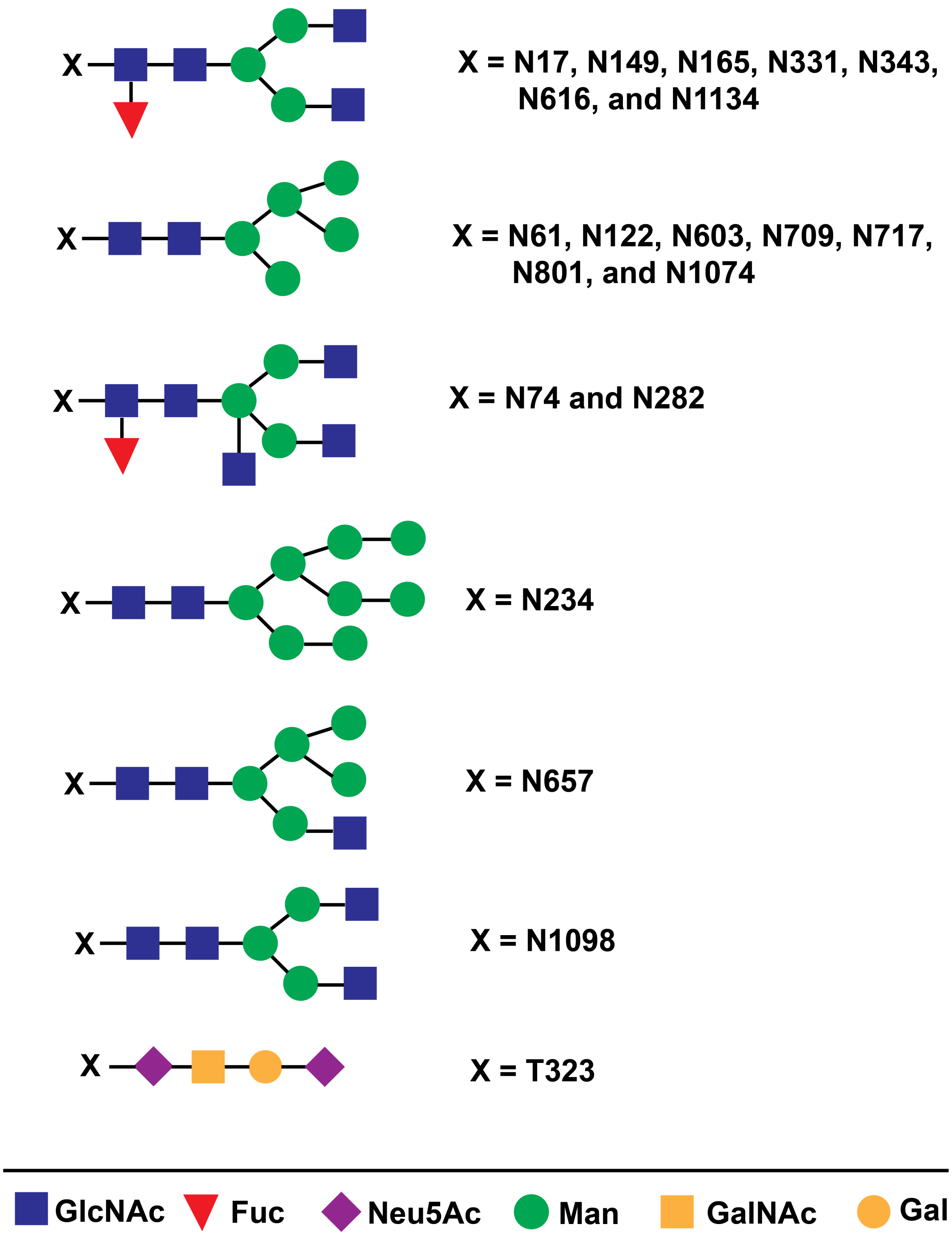


**Figure S13.** Glycan compositions used in this study.

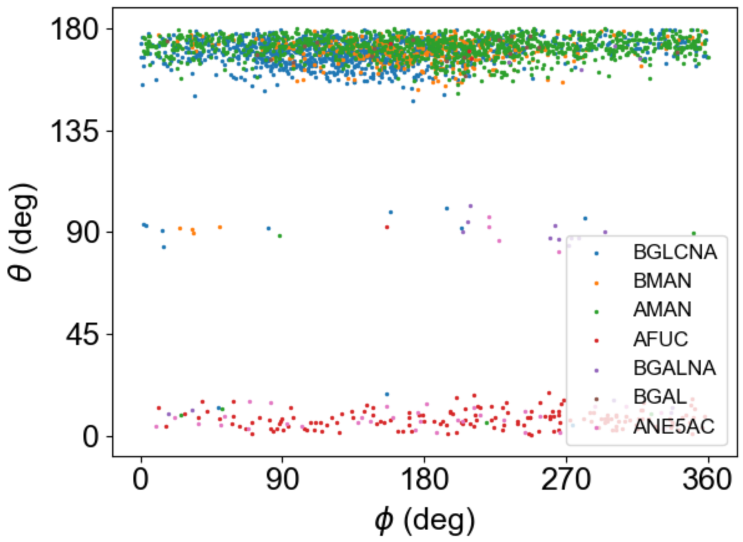


**Figure S14.** Cremer-Pople angles[13] of the glycans after simulated annealing.

**Table S3.** Details of the antibodies investigated in this study. Antibodies 2-4[14], 4A8[15], and BD-368-2[16] were obtained from B cells of patients after SARS-CoV-2 infection, while S309[17] was extracted from B cells of SARS-CoV infected patients. Antibodies CR3022[18] and STE90-C11[19] were created from phage libraries.

| Antibody | Binds to | Residues in epitope | PDB id | Reference |
|---|---|---|---|---|
| 2-4 | SARS-CoV-2 RBD | 446–447, 449, 452–453, 455–456, 483–487, 489, 490, 492–496, 498 | 6XEY | Liu et al.[14] |
| 4A8 | SARS-CoV-2 NTD | 143–148, 150–152, 158, 245–251, 256–257 | 7C2L | Chi et al.[15] |
| BD-368-2 | SARS-CoV-2 RBD | 346, 351, 444–447, 449–450, 452, 470, 472, 478–486, 490, 492, 494, 498 | 7CHH | Du et al.[16] |
| CR3022 | SARS-CoV-2 RBD | 369–372, 374–386, 389–390, 392, 427–430, 515–517, 519 | 6W41 | Yuan et al.[18] |
| S309 | SARS-CoV/SARS-CoV-2 RBDs | 333–335, 337, 339–341, 343–346, 354, 356–361, 440, 441, 444, 509 | 6WPS | Pinto et al.[17] |
| STE90-C11 | SARS-CoV-2 RBD | 403, 405, 406, 408, 409, 415–417, 420, 421, 449, 453, 455–460, 473–477, 486–487, 489, 493–496, 498, 500–505 | 7B3O | Bertoglio et al.[19] |
| S2M11 | SARS-CoV-2 RBD | Up protomer: 339, 342, 343, 345, 367, 368, 371–374, 436, 440, 441, 444, 339, 342, 343, 345, 367, 368, 371–374, 436, 440, 441, 444<br>Down protomer: 446, 447, 449, 452, 455, 456, 483–490, 492–494, 496, 498 | 7K43 | Tortorici et al.[20] |
